# Cell cycle-controlled clearance of the CcrM DNA methyltransferase by Lon is dependent on DNA-facilitated proteolysis and substrate polar sequestration

**DOI:** 10.1101/293738

**Authors:** Xiaofeng Zhou, Lucy Shapiro

## Abstract

N6-adenine methylation catalyzed by the DNA methyltransferase CcrM is an essential epigenetic event of the *Caulobacter* cell cycle. Limiting CcrM to a specific time period during the cell cycle relies on temporal control of *ccrM* transcription and CcrM proteolysis. We investigated how Lon, a protease from AAA+ superfamily conserved from bacteria to humans, temporally degrades CcrM to maintain differential chromosomal methylation state, thereby regulating transcription factor synthesis and enabling cell cycle progression. We demonstrate that CcrM degradation by Lon requires DNA as an adaptor for robust proteolysis. Lon, a DNA-bound protein, is constitutively active throughout the cell cycle, but allows CcrM mediated DNA methylation only when CcrM is transcribed and translated upon completion of DNA replication. An additional mechanism to limit CcrM activity to a narrow window of the cell cycle is its sequestration to the pole of the progeny stalked cell, which prevents physical contact with DNA-bound Lon. Thus, we have provided evidence for a novel mechanism for substrate selection by the Lon protease, providing robust cell cycle control mediated by DNA methylation.

## Introduction

Epigenetic regulation of gene expression by DNA methylation is a conserved mechanism in all domains of life (Casadesús and Low, 2006; He et al., 2011; Smith and Meissner, 2013). In most mammalian and plant cells, DNA methylation refers to the addition of a methyl group to the cytosine bases in the contexts of CG, CHG, and CHH (H = A, C, or T) (Kim and Zilberman, 2014; Lister et al., 2009). In bacteria, DNA methylation was originally discovered as a component of restriction-modification (R-M) systems consisting of an endonuclease and an associated DNA methyltransferase, used to differentiate the genome DNA from invading phage DNA (Bickle and Krüger, 1993). However, several solitary DNA methyltransferases without apparent cognate restriction enzymes were later identified in many bacterial genomes (Collier, 2009; Sánchez-Romero et al., 2015). These orphan N6-adenine DNA methyltransferases were found to regulate the initiation of chromosome replication, DNA mismatch repair, gene expression, and cell cycle progression (Collier, 2009; Gonzalez et al., 2014; Iyer et al., 2006; Reisenauer et al., 1999; Val et al., 2012). The two best-studied examples are the *Escherichia coli* Dam enzyme (methylating the adenine of GATC) and the *Caulobacter crescentus* CcrM enzyme (methylating the adenine of GANTC).

The α-proteobacterium *Caulobacter crescentus* is a model system for elucidating the mechanisms leading to an asymmetric cell division. *Caulobacter* produces two morphologically distinct progeny at each cell division: a motile swarmer progeny (SWP) and a sessile stalked progeny (STP) (Figure 1A). The progeny swarmer cell (G1 phase) cannot initiate chromosome replication until it differentiates into a stalked cell (ST), whereas the progeny stalked cell immediately initiates chromosome replication and enters S phase (Figure 1A and 1B). *Caulobacter* initiates chromosome replication once and only once per cell cycle (Marczynski and Shapiro, 2002). Replication initiates on a fully methylated chromosome (adenine of GANTC sites is methylated on both strands) and the movement of the replication fork culminates in the generation of two hemi-methylated chromosomes (adenine of GANTC is methylated on only one of the two strands) (Figure 1B) (Kozdon et al., 2013). Upon completion of chromosome replication in the pre-divisional (PD) cell (Figure 1A and 1B), a burst of CcrM protein synthesis converts the hemi-methylated chromosomes back into two fully methylated chromosomes (Figure 1B). The methylation state of GANTC motifs within a subset of promoters directly regulates the transcription of genes comprising the cyclical genetic circuit that drives the cell cycle (Figure 1C). DnaA serves as an initiator of chromosome replication and as a transcription factor that controls the transcription of approximately 50 cell cycle-regulated genes (Hottes et al., 2005). Efficient transcription of *dnaA* (located close to the origin of replication) requires the GANTC site within its promotor to be in the fully methylated state. Upon replication initiation, the passage of the replication fork converts the *dnaA* promoter from the fully methylated state to the hemi-methylated state, thus turning down the transcription of *dnaA* (Collier et al., 2007). As replication proceeds, the *ctrA* gene, which is positioned further from the replication origin, transitions from the fully methylated state to the hemi-methylated state. In the case of the *ctrA* promoter, it is activated when in the hemi-methylated state (Reisenauer and Shapiro, 2002). The transcription of *ctrA* is controlled by two promoters, one of which, the *ctrA*P1, is regulated by DNA methylation in a GcrA-dependent manner. The *ctrA*P1 is activated after the replication fork passes through the *ctrA*P1 and it becomes hemi-methylated. GcrA stabilizes RNA polymerase holoenzyme by interacting with σ70, and stimulates open complex formation via the presence of a preferred methylation site near the *ctrA*P1 (Haakonsen et al., 2015). DnaA and CtrA have opposite modes of function and regulation of their transcription. DnaA activates initiation of DNA replication, and transcription of the *dnaA* gene is activated when its promoter is in a fully methylated state. CtrA inhibits initiation of DNA replication, and its transcription is activated by GcrA when its promoter is in the hemi-methylated state. The methylation state of the chromosome, which is temporally modulated by the passage of the replication fork, controls the sequential expression of the DnaA and CtrA master transcription factors, which provide a regulatory hierarchy that activates or represses >300 cell cycle-regulated genes (Zhou et al., 2015).

**Figure 1.**
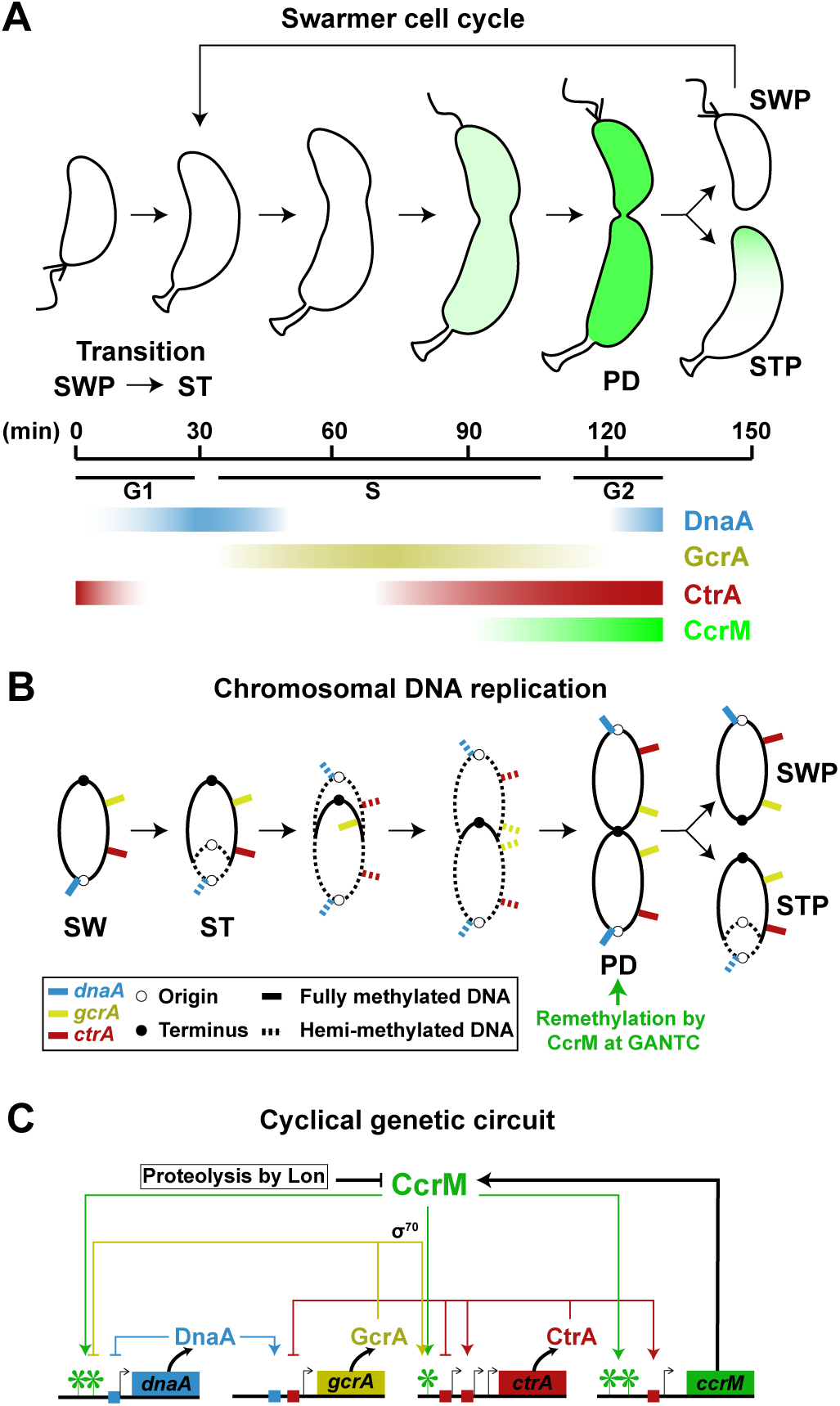
CcrM-mediated DNA methylation regulates the cell cycle control circuit by linking the progression of the cell cycle to chromosome replication. (A) Schematic of the *Caulobacter* cell cycle. Stages of the *Caulobacter* cell cycle are shown in 30 min intervals, beginning with the swarmer progeny (SWP) at 0 min. Swarmer cells (G1 phase) develop into stalked cells (ST) and enter an S phase. As stalked cells elongate and become pre-divisional cells (PD), CcrM (green) is synthesized. The pre-divisional cells begin compartmentalization (G2 phase), yielding two morphologically distinct daughter swarmer (SWP) and stalked (STP) cells. The temporal distribution of global regulators, DnaA (blue), GcrA (yellow), and CtrA (red), are shown. (B) Schematic showing the changes in the methylation state of GANTC motifs on the chromosome as a function of chromosome replication and cell cycle progression. The locus of *dnaA* (blue), *gcrA* (yellow), *ctrA* (red), and *ccrM* (green) and their corresponding methylation states are indicated. (C) CcrM plays a central role in the regulation of a cyclical genetic circuit driving *Caulobacter* cell cycle. The asterisk indicates fully methylated GANTC site.

CcrM is present only during a narrow window of the cell cycle (Figure 1A) coincident with its time of transcription and translation (Schrader et al., 2016; Zhou et al., 2015). Following its burst of synthesis, CcrM is cleared from the cell by the Lon protease (Wright et al., 1996). Lon is a member of the AAA+ protease superfamily that is widely distributed in all kingdoms of life. In a *lon* deficient strain, CcrM remains detectable throughout the cell cycle, leading to the accumulation of multiple chromosomes (Wright et al., 1996; Zweiger et al., 1994). Constitutive overexpression of CcrM results in mis-regulation of over 10% of cell cycle-controlled genes due to the aberrant GANTC methylation state of their promotors (Gonzalez et al., 2014), demonstrating that restricted presence of CcrM to a short time window is essential for controlling cell cycle progression. Because Lon is normally present throughout the cell cycle, the pre-divisional cell undergoes an “arms race” between CcrM synthesis and degradation (Wright et al., 1996). The mechanism that protects CcrM from Lon-mediated degradation in the pre-divisional cell has remained enigmatic.

In this study, we determined that the robust degradation of CcrM requires the presence of DNA as an adaptor. Lon-mediated proteolysis of CcrM occurs when CcrM binds DNA eliciting a race between catalysis of N6-adenine methylation and CcrM degradation. The affinities of CcrM-DNA and Lon-DNA are 10-fold higher than that of the direct interaction between CcrM and Lon. High levels of newly synthesized CcrM in the pre-divisional cell tilts the race towards complete methylation of ~4500 chromosomal GANTC sites. Upon cell division, CcrM synthesis stops and Lon degradation of the remaining CcrM wins the race. Each progeny cell acquires a single fully methylated chromosome. The daughter swarmer cell, which cannot initiate DNA replication, exhibits complete degradation of CcrM, in part due to the extended time span of the swarmer-to-stalked cell transition. However, the progeny stalked cell has inadequate time for the complete degradation of remaining CcrM before its immediate initiation of DNA replication. This raises the problem of how the hemi-methylated state of newly synthesized chromosomal DNA is maintained in the stalked progeny during the ensuing replication fork progression. We show here that excess CcrM is sequestered to the pole of the cell away from the chromosome while Lon is bound to DNA, allowing the propagation of hemi-methylated DNA during replication. The sequestration of remaining CcrM begins with the initiation of chromosome replication and ends prior to the formation of the division plane. The robust combination of CcrM sequestration and clearance of DNA-bound CcrM by Lon protects the replicating chromosome from re-methylation, thereby coordinating gene expression and replication fork progression.

## Results

### The C-terminal domain of CcrM is required for degradation by the Lon protease and for methyltransferase activity

ATP-dependent proteases usually rely on terminal sequences for substrate recognition (Joshi and Chien, 2016; Sauer and Baker, 2011). To identify the CcrM degradation tag, we fused an M2 epitope to the N- or C-terminus of natively expressed CcrM on the chromosome. Swarmer cells expressing a sole copy of M2-CcrM or CcrM-M2 were isolated and allowed to proceed synchronously through the cell cycle. Samples were collected every 20 minutes for immunoblot analysis using an anti-CcrM antibody. We observed that M2-CcrM was proteolyzed until the completion of DNA replication, whereas CcrM-M2 was present throughout the cell cycle, indicating that the C-terminal M2 tag protected CcrM from degradation by interfering with its recognition by Lon (Figure 2A).

**Figure 2.**
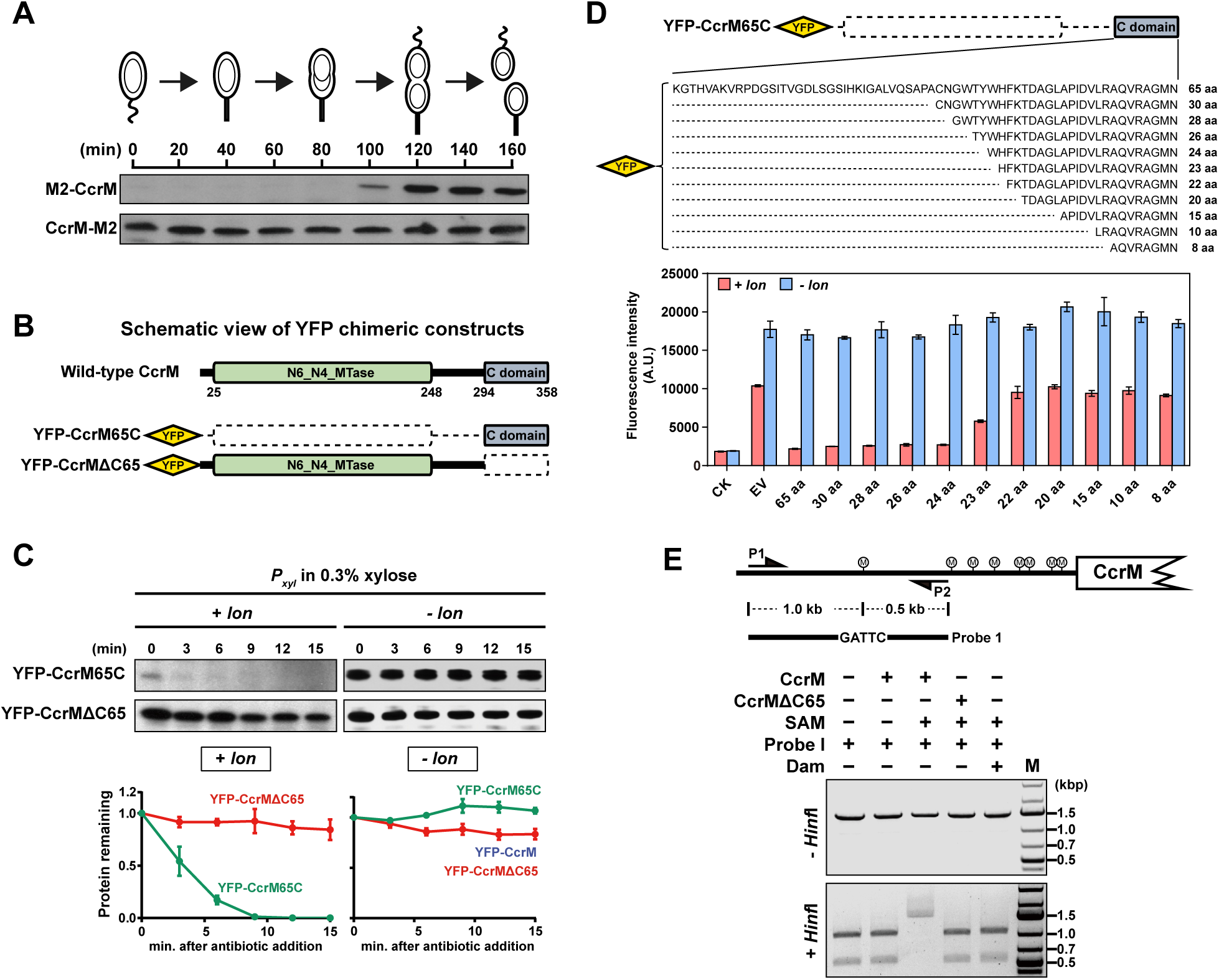
The C-terminus of CcrM is necessary for recognition by Lon and for DNA methyltransferase activity. (A) Cells containing a single copy of M2-CcrM or CcrM-M2 under the control of native promotor were grown in M2G, synchronized, and released onto fresh M2G. Samples were taken every 20 min for immunoblots with anti-CcrM antibody. (B) Schematic representation of CcrM domain structure and YFP chimeric constructs used in this study. Dash line indicates a deletion of amino acids. The numbers refer to amino acid positions. (C) *In vivo* degradation assays showing the effect of C-terminal 65 residues on CcrM protein stability. Stabilities of YFP chimeric proteins in ∆*lon* (*- lon*) cells are shown for comparison. Cells were grown in PYE with 0.3% xylose to exponential phase and treated with antibiotics for protein synthesis shut-off assays. Protein levels were monitored by immunoblot using anti-GFP antibody (top). Band intensities were quantified (bottom) and error bars represent SDs (*n* = 3). (D) Florescence of wild-type (*+ lon*) and ∆*lon* (*- lon*) harboring plasmids expressing chimeric YFP proteins. A schematic shows a series of truncations in which 8, 10, 15, 20, 22, 23, 24, 26, 28, and 30 amino acids are retained from CcrM C-terminal 65 amino acids (top). The florescence normalized by optical density is shown (bottom). The means ± SDs (*n* = 3) are plotted. CK, cells expressing free YFP. EV, cells expressing YFP-CcrM65C. (E) DNA methylation assay showing the effect of CcrM C-terminal domain on its DNA methyltransferase activity. A schematic of Probe 1 designing strategy is shown (top). PCR amplified Probe 1 was incubated with CcrM or CcrMΔC65 in the presence of S-adenosyl methionine (SAM). DNA methylation states were assayed by *Hin*fI digestion (bottom). Dam methylase from *E. coli* served as a negative control. P1 and P2 are primers for Probe 1 amplification.

To further validate Lon recognition of the C-terminus of CcrM, truncations lacking the N-terminal 294 residues of CcrM (CcrM65C) or the C-terminal 65 amino acids (CcrMΔC65) were generated and fused to YFP (Figure 2B). The *in vivo* degradation rates of these chimeric proteins were measured in wild-type cells and in cells bearing a deletion of Lon. In the presence of Lon, YFP-CcrM65C was extremely unstable and had a half-life of ~ 3 min, whereas YFP-CcrMΔC65 was stable (Figure 2C). In the absence of Lon, both YFP-CcrM65C and YFP-CcrMΔC65 were all stable (Figure 2C). Our data indicate that the CcrM C-terminus is necessary and sufficient for Lon recognition.

To determine the precise amino acid sequence of the CcrM degradation tag within the C-terminal 65 amino acids, we generated a series of truncations based on the YFP-CcrM65C construct (Figure 2D). Turnover of the chimeric proteins was quantified using a fluorescent microplate reader. The fluorescence levels of wild-type *Caulobacter* strains harboring plasmids expressing YFP chimeric protein containing 24, 26, 28, 30, or 65 C-terminal amino acids from CcrM were all significantly depressed, suggesting that the C-terminal 24 amino acids of CcrM are sufficient to confer Lon-dependent proteolysis (Figure 2D). The fluorescence data reflect the stability of the chimeric proteins as all YFP chimeric proteins expressed in a Lon deletion strain maintained high levels of fluorescence.

Given that the CcrM C-terminus is required for its proteolysis, we asked whether the 65 amino acids within the CcrM C-terminus are also required for enzymatic function. We performed *in vitro* DNA methylation assays using purified CcrM and CcrMΔC65 in the presence of a DNA fragment (hereafter named Probe 1) that contains one methylation site (GATTC) (Figure 2E). The distances from the methylation site to 5’ and 3’ ends of the probe were 1.0 kb and 0.5 kb, respectively. The restriction enzyme *Hinf*I can be used to distinguish methylated and unmethylated DNA because it cuts only the unmethylated GANTC sequence. When incubated with CcrM and the methyl group donor S-adenosyl methionine (SAM), *Hinf*I was unable to digest Probe 1, indicating that the GATTC site was methylated (Figure 2E). In contrast, Probe 1 incubated with CcrMΔC65 was digested by *Hinf*I, giving two fragments at 1.0 kb and 0.5 kb in length on agarose gels. Combined, our results demonstrate that C-terminus of CcrM is required for both DNA-methyltransferase activity and proteolysis by Lon.

### Conserved C-terminal motifs determine CcrM DNA binding activity

Sequence alignment of the CcrM C-terminus revealed four highly conserved motifs among CcrM homologues in α-proteobacteria (Figure S1A). To investigate the roles of C-terminal conserved motifs in CcrM function, we generated CcrM mutants using alanine substitution at a conserved residue within each motif (Figure S1A). Mutations of *ccrM* bearing the alanine-substitutions shown in Fig S1A were introduced into a *ccrM* depletion strain and expressed under the control of the native *ccrM* promotor. In the absence of wild-type CcrM, mutations at S315 and W332 caused severe defects in viability, cell division and morphology, exhibiting filamentous bacterial growth due to the essentiality of CcrM protein function (Figure S1B-S1D). *In vitro* gel shift assays were performed using purified CcrM and CcrMS315A to test their DNA binding activity. The results revealed that wild type CcrM bound Probe 1, but CcrMS315A did not, indicating that the S315 mutation abolished DNA binding activity (Figure S1E). A W332A mutation was shown to lack DNA binding activity (Dr. Norbert O. Reich, personal communication). Thus, two motifs within the conserved C-terminus of CcrM are required for DNA binding activity.

### Lon protease binds to DNA and is constitutively active during the *Caulobacter* cell cycle

We previously showed that Lon protein abundance does not change during *Caulobacter* cell cycle (Wright et al., 1996). Although the protein level of Lon is constant, it is possible that Lon activity is cell cycle-dependent. To address this possibility, we assayed Lon activity as a function of cell cycle progression using a known substrate that is degraded directly by Lon in the absence of adaptors and any accessory factors. To circumvent the possibility that the substrate to be tested has cell cycle-dependent regulation, we generated an exogenous Lon substrate by tagging the C-terminus of the YFP protein with a sul20C Lon degradation tag (Gur and Sauer, 2009) driven by the P_*xyl*_ promoter with its constitutive expression induced by xylose. We observed that YFP-sul20C was degraded by Lon in a wild-type background, but not in a *Δlon Caulobacter* mutant (Figure 3A). The YFP-sul20C substrate was used to assess Lon proteolytic activity in swarmer, stalked, and pre-divisional cells obtained from synchronized cultures. The results demonstrate that the degradation rate of YFP-sul20C is not significantly different in three types of cells, suggesting that Lon activity is cell cycle-independent (Figure 3B).

**Figure 3.**
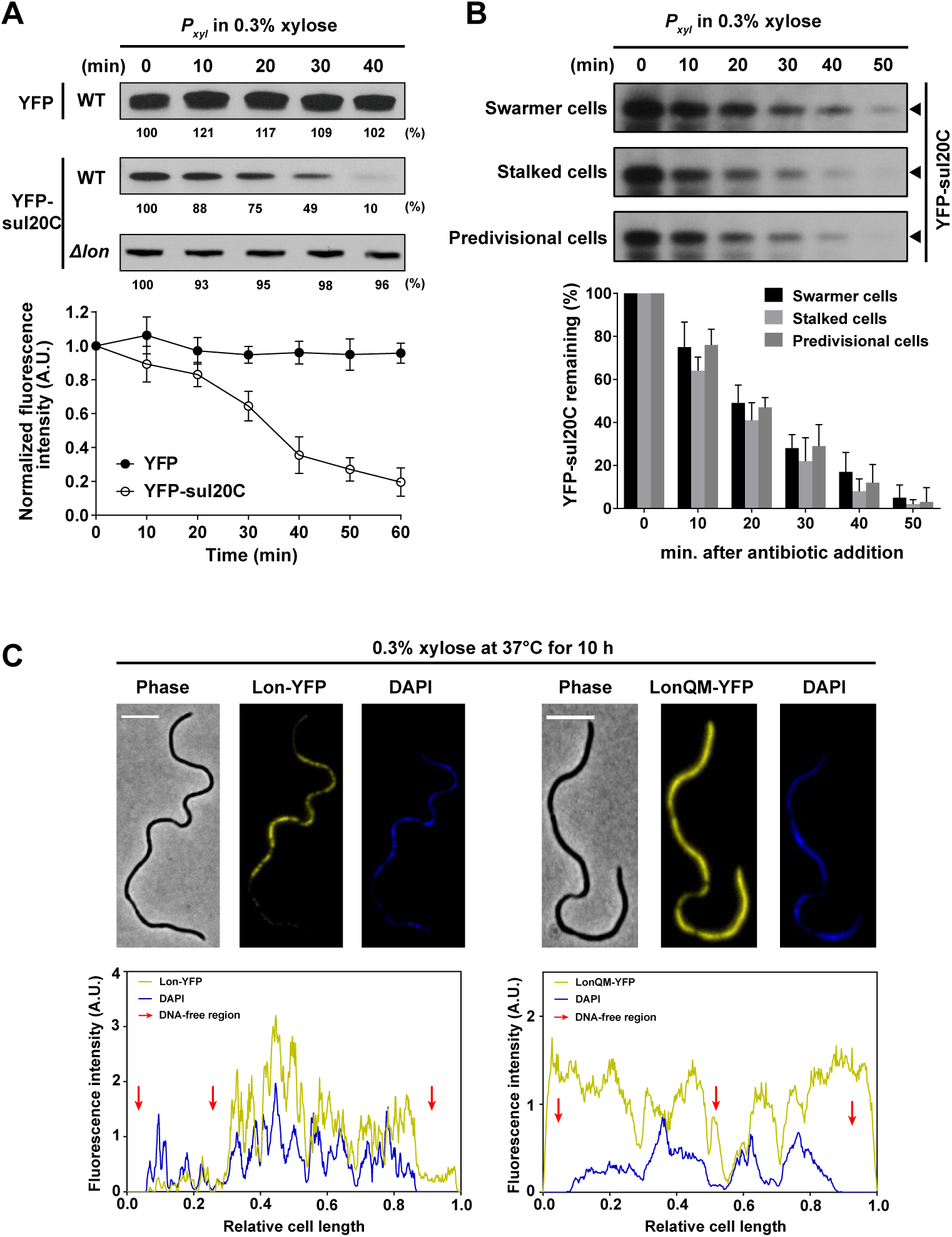
Lon is a DNA-binding protein and its proteolytic activity is constitutively active during *Caulobacter* cell cycle. (A) *In vivo* degradation assays showing stabilities of YFP and YFP-sul20C in wild-type. YFP-sul20C stability in ∆*lon* cells is shown for comparison. Merodiploid cells expressing free YFP or YFP-sul20C were grown in PYE with 0.3% xylose to exponential phase and treated with antibiotics for protein synthesis shut-off assays. Protein levels were monitored by immunoblot using anti-GFP antibody (top). Band intensities were quantified and indicated as percentage. The cellular florescent intensity normalized by optical density is measured (bottom). The means ± SDs (*n* = 4) are plotted. (B) *In vivo* degradation assays showing YFP-sul20C stabilities in swarmer, stalked, and pre-divisional cell. Cells expressing YFP-sul20C controlled by *P*_*xyl*_ were grown in M2G with 0.3% xylose, synchronized, and harvested at 0, 60, and 120 mps. Samples were treated with antibiotics for protein synthesis shut-off assays. Protein levels were monitored by immunoblot using anti-GFP antibody (top). Band intensities were quantified (bottom) and error bars represent SDs (*n* = 3). (C) Fluorescence images showing Lon-YFP colocalizing with DAPI-stained DNA in a *Caulobacter* temperature-sensitive *parE* and *ftsA* mutant (PC6340) that produces DNA-free regions. LonQM-YFP lacking DNA binding activity is shown for comparison. Cells were cultured at the restrictive temperature (37^°^C) for 10h in M2G medium with 0.3% xylose prior to DAPI staining and imaging (top). Scale bar = 5 µm. Fluorescence intensity profiles of Lon-YFP or LonQM-YFP and DAPI signals along the long axis of the cell are shown (bottom). Red arrows indicate DNA-free regions.

In *E. coli*, up to 95% of Lon molecules are bound to DNA (Karlowicz et al., 2017). To determine if Lon co-localizes with DNA in *Caulobacter*, we integrated a plasmid bearing a translational fusion of YFP to the C-terminus of Lon under the control of P_*xyl*_ into the chromosome of a temperature-sensitive (*ts*) mutant that forms filamentous cells when grown at the restrictive temperature, generating large DNA-free regions (Ward and Newton, 1997). A similar construct was made using a Lon mutant (LonQM) that lacks DNA binding activity (Figure S2A). We ruled out any effect of fusing YFP to Lon N- or C-terminus on Lon function (Figure S2B). Cultures of the *ts* mutant bearing the Lon-YFP construct were grown at restrictive temperature in the presence of xylose and imaged by epifluorescence microscopy. We observed that Lon-YFP co-localized with the DAPI DNA signal and was absent in DNA-free regions (Figure 3C). In contrast, the LonQM-YFP signal was observed throughout the entire cell, including the DNA-free regions (Figure 3C). Cumulatively, these results suggest that Lon binds DNA *in vivo* and its proteolytic activity is cell cycle-independent.

### Both Lon and CcrM are capable of binding to the same DNA probes with high affinity

DNA Binding by CcrM is a prerequisite for its chromosome methylation activity. Given that DNA-bound Lon protease is active throughout the cell cycle, we hypothesized that CcrM, which has been shown to processively move on DNA (Berdis et al., 1998; Woodcock et al., 2017), could be recognized by Lon bound to DNA. To test this hypothesis, we attempted to reconstitute this interaction *in vitro* using three different DNA probes. Besides the Probe 1 used in our previous methylation assays (Figure 2E), we designed Probe 2 by mutating Probe 1’s methylation site from GATTC to AATAC. Probe 3 is from a region upstream of the *pilA* gene lacking any GANTC motif (Figure 4A). Gel shift assays demonstrated that the purified CcrM protein can bind Probes 1, 2, and 3, suggesting that the DNA binding capability of CcrM does not require the GANTC motif (Figure 4B). As expected, in this assay purified CcrMΔC65, lacking the DNA binding domain failed to exhibit DNA binding activity (Figure 4B). This results also accounts for the observations that the CcrM mutations at S315 and W332 led to complete inactivation of DNA binding and methyltransferase activity (Figure S1). We found that the purified Lon protease also binds to all three probes (Figure 4B). Because unmethylated DNA is absent *in vivo*, we sought to investigate the DNA binding capabilities of CcrM and Lon using hemi-methylated and fully-methylated DNA probes. We obtained fully-methylated DNA by incubating PCR-generated Probe 1 with purified CcrM protein. The hemi-methylated DNA probe was generated by hybridization of fully-methylated and unmethylated DNA probes. The methylation states of these DNA probes were confirmed by overlapping restriction digestions (Figure S3A and S3B). We found that the binding capabilities of CcrM to DNA, as well as to the Lon protease, are methylation state-independent (Figure S3C). Our results support the hypothesis that both CcrM and Lon are capable of binding to DNA probes simultaneously and independent of methylation state.

**Figure 4.**
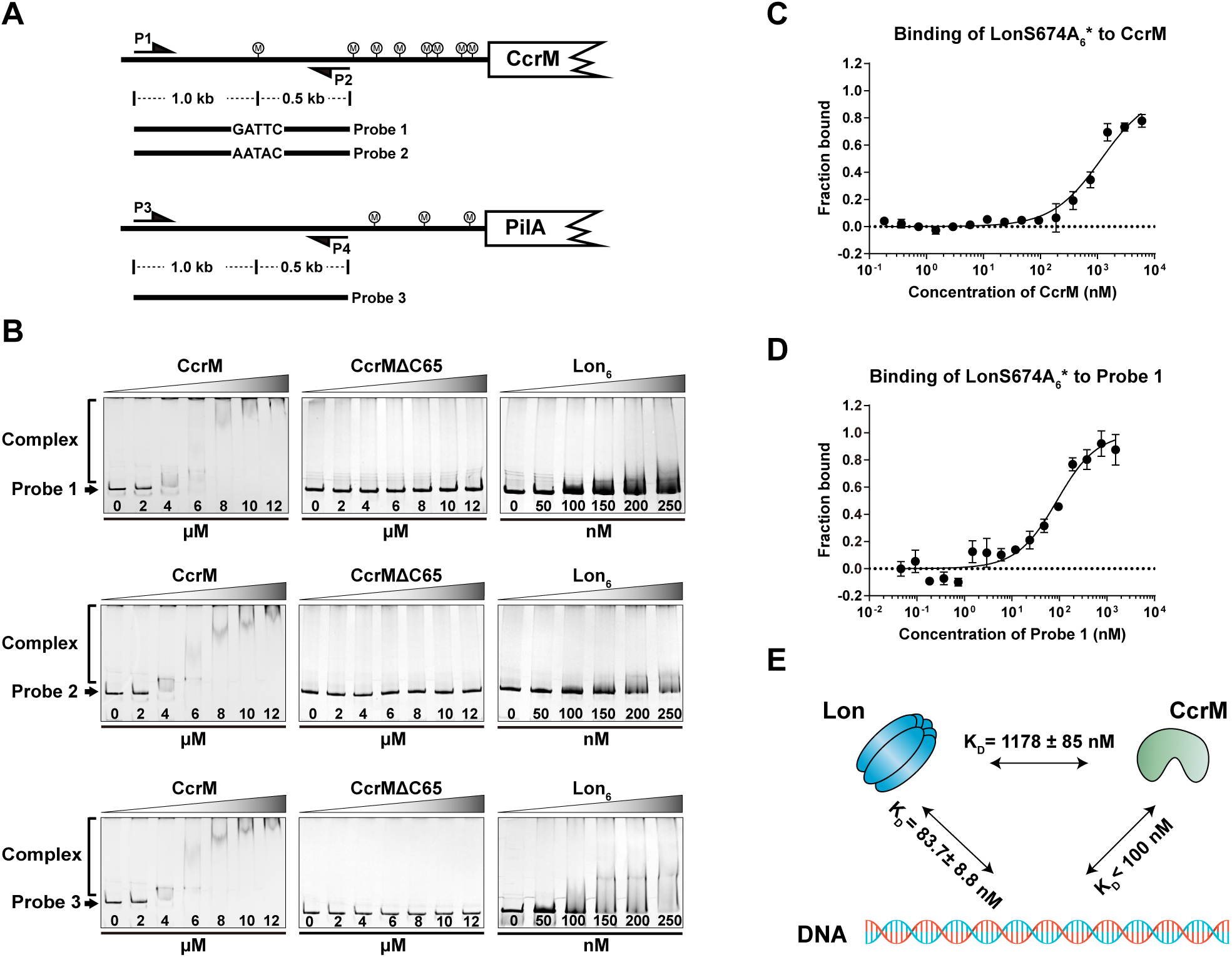
CcrM binds DNA probes *in vitro* with high affinity. (A) Schematic view of DNA probe designs according to genome locus. Probe 2 is designed based on Probe 1 with mutation of GATTC to AATAC. Probe 3 is designed from the upstream sequence of *pliA*. P1-P2 and P3-P4 are primers to amplify Probe 1 or 2 and Probe 3, respectively. CcrM methylation sites are shown. (B) EMSA showing binding of recombinant CcrM, CcrMΔC65, or Lon to Probe 1, 2, and 3, respectively. See Methods for experimental details. (C-D) The direct binding of purified LonS674A to CcrM or Probe 1 was assessed *in vitro* by microscale thermophoresis. LonS674A was fluorescently labeled with Atto-488 dye, indicated by LonS674A^*^. The concentration of LonS674A_6_^*^ was held constant at 20 nM while CcrM (C) or Probe 1 (D) was titrated in 2-fold serial dilutions against it. The purified proteins were allowed to incubate together at room temperature for 10 min prior to the binding assay. The data report the fraction of LonS674A_6_^*^ that is bound at each concentration of CcrM (C) or Probe 1 (D). See Methods for description of curve fits. (E) Cartoon depicting affinities measured in Figure 4C and 4D between CcrM, Lon, and DNA. CcrM and Lon have affinities to DNA ~14 folds more than that of CcrM-Lon direct interaction.

To measure the affinities of Lon binding to DNA and Lon binding to CcrM, we used microscale thermophoresis (MST) assays (Wienken et al., 2010). To perform MST assays, we first labeled lysine residues on LonS674A, a mutant protein that lacks proteolytic activity but retains DNA binding activity (Figure S2A) (Botos et al., 2004; Karlowicz et al., 2017), with the Atto-488 dye, as indicated by LonS674A^*^. We then measured the change in the thermophoresis of LonS674A^*^ over a 2-fold serial dilution of either CcrM or Probe 1. Direct binding was observed between LonS674A^*^ and CcrM (*K*_*D*_ = 1178 ± 85 nM) (Figure 4C) and between LonS674A^*^ and Probe 1 (*K*_*D*_ = 83.7 ± 8.8 nM) (Figure 4D). Thus, there is a ~14-fold-weaker affinity between LonS674A^*^ and CcrM than between LonS674A^*^ and DNA Probe 1. Recent studies on DNA recognition by CcrM reported an equilibrium dissociation constant of 108 ± 20 nM for double-stranded DNA (Woodcock et al., 2017). We performed a quantitative Western blot to determine the concentration of CcrM *in vivo* at the 120 minutes-post-synchrony timepoint, using purified CcrM to calibrate a standard curve (Figure S3D). We determined that the intracellular concentration of CcrM ranged from 950 - 1280 nM over three measurements, averaging 1090 ± 135 nM for pre-divisional cells (Figure S3D). The highest intracellular concentration of CcrM approached the *K*_*D*_ value of CcrM-Lon direct interaction. These findings demonstrate that both Lon and CcrM associates with DNA in vivo and in vitro, suggesting that degradation of CcrM in vivo may occur while it is bound to DNA (Figure 4E).

### DNA plays an adapter role in CcrM proteolysis by Lon

To test whether the presence of DNA can stimulate CcrM proteolysis by Lon, we performed *in vitro* degradation assays in the presence of the DNA probes described in Figure 4A. The addition of Probe 1, containing the GATTC methylation recognition site, dramatically boosted CcrM degradation (Figure 5A). Strikingly, the addition of Probe 2 (the same as Probe 1 but with a scrambled DNA methylation site) or Probe 3 (with a non-specific DNA sequence) produced CcrM degradation rates similar to that observed in the presence of Probe 1. These results suggest that CcrM degradation stimulated by DNA does not depend on the presence of a methylation site or a specific DNA sequence (Figure 5A). As a negative control, the degradation of CcrMΔC65 was not observed in the presence or absence of DNA (Figure 5A). Titration of DNA showed that increasing concentrations increased the rate of CcrM proteolysis, but reached a maximum rate of degradation around 10 nM concentration of DNA (Figure 5B). DNA has been reported to stimulate Lon ATPase activity (Charette et al., 1984; Chung and Goldberg, 1982; Zehnbauer et al., 1981). We found that both DNA and the degradation substrate CcrM stimulate Lon ATPase activity, but that addition of both did not further stimulate the ATPase (Figure 5C). Degradation kinetics of β-casein, a non-DNA binding Lon substrate, was also measured in the presence and absence of DNA. We did not observe stimulated β-casein proteolysis, indicating that DNA-facilitated proteolysis might be restricted to DNA-binding substrates (Figure S4A). To test whether stimulated proteolysis requires both CcrM and Lon to bind DNA, we performed *in vitro* degradation assays using previously identified DNA-binding deficient mutants, CcrMS315A (Figure S1E) and LonQM (Figure S2A). The LonQM mutant exhibited intact proteolytic activity on β- casein, a non-DNA binding substrate (Figure S4B). We found that DNA failed to stimulate CcrMS315A degradation by wild-type Lon. Similarly, CcrM degradation by LonQM was not stimulated by the addition of DNA (Figure 5D). Thus, DNA-facilitated proteolysis requires both protease and substrate to bind DNA.

**Figure 5.**
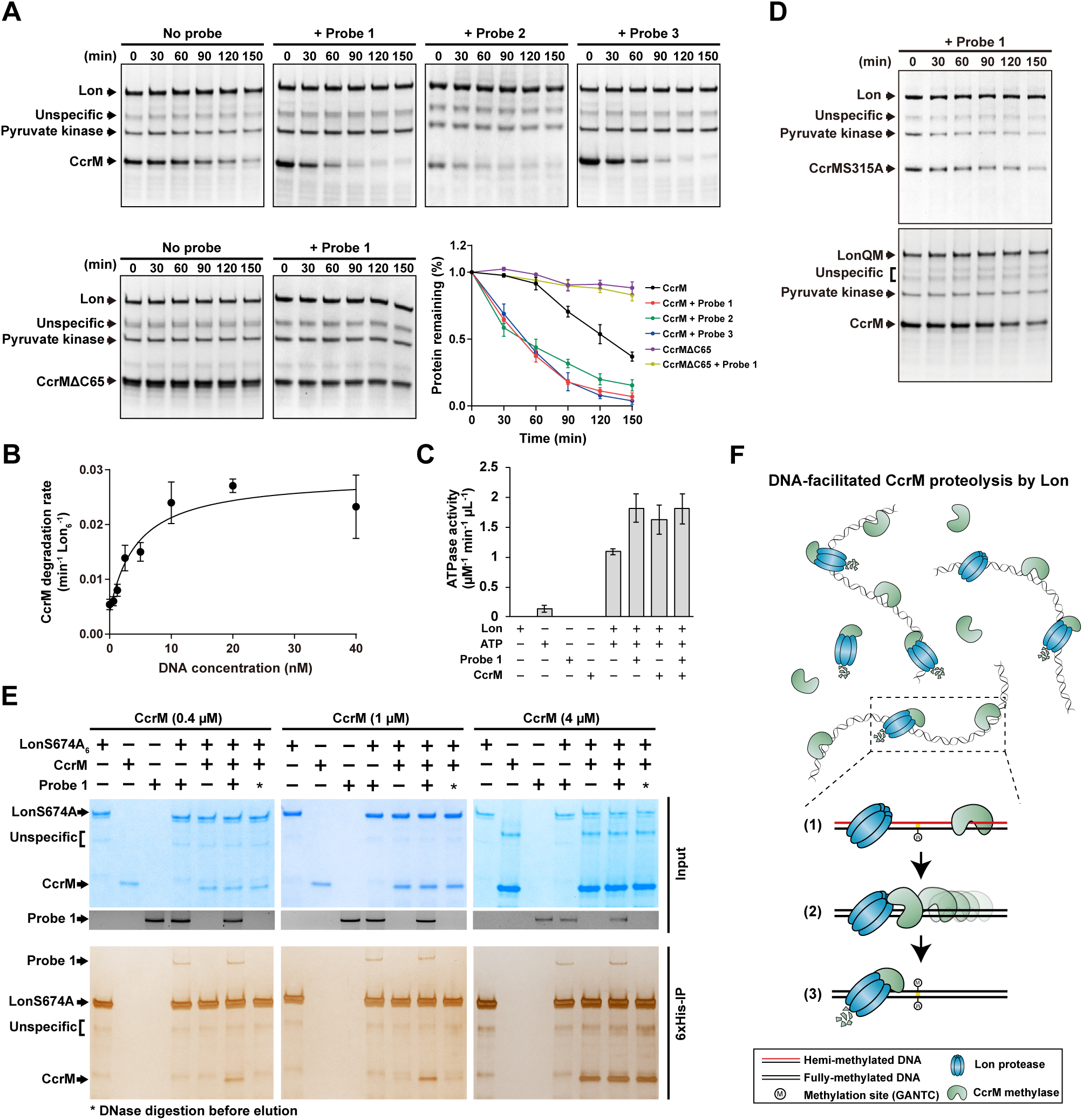
DNA serves as an adaptor for Lon-mediated CcrM proteolysis. (A) *In vitro* degradation assays showing the stimulatory effect of DNA on CcrM degradation by Lon. CcrM (1 µM) was incubated with Lon_6_ (0.2 µM) in the absence or presence of DNA probes (10 nM). Degradation of CcrMΔC65 was also assayed in the presence or absence of DNA probe as indicated. The intensity of CcrM or CcrMΔC65 bands from three independent experiments were quantified and plotted. (B) DNA-facilitated CcrM degradation by Lon. Degradation rates of CcrM (1 µM) by Lon (0.2 µM) are shown for increasing concentration of DNA probe. See Methods for description of curve fits. (C) ATPase activity of Lon in presence and absence of DNA and CcrM. See Methods for detailed description of ATPase assay. (D) *In vitro* degradation assays showing the degradation of CcrMS315A by Lon and the degradation of CcrM by LonQM. CcrMS315A or CcrM (1 µM) was incubated with Lon or LonQM_6_ (0.2 µM) in the absence of Probe 1 (10 nM). Pyruvate kinase is part of the ATP regeneration system. (E) DNA facilitates recognition of CcrM by LonS674A in a low concentration. Coomassie-stained SDS-PAGE gels showing co-immunoprecipitation of nucleoprotein complex. The concentration of LonS674A_6_ was maintained at 0.2 µM. A low concentration of CcrM (0.4 µM) requires the presence of DNA to be recognized by LonS674A (left), whereas the recognition of a high concentration of CcrM (4 µM) does not depend on the presence of DNA (right). Asterisks indicate DNase I digestion before elution. (F) Cartoon depicting DNA-facilitated CcrM degradation by Lon. The left panel shows the presence of CcrM, Lon, and DNA fragments in a mixed reaction. A zoomed-in schematic view (right panel) shows the three steps of CcrM degradation by Lon on DNA: (1) preferential binding of CcrM and Lon to DNA fragments due to their individual high affinity; (2) enhanced-intermolecular collision frequency driven by CcrM processivity; (3) substrate unfolding and proteolysis. DNA plays dual roles in modulating CcrM-mediated adenine methylation and CcrM degradation by Lon.

Further, we performed co-immunoprecipitation (Co-IP) of the reconstituted reactions to determine whether CcrM-Lon-DNA can form nucleoprotein complexes. We first conducted Co-IP using low concentrations of CcrM substrate (0.4 µM). CcrM co-immunoprecipitated with LonS674A only if DNA was present, demonstrating that recognition of CcrM by Lon relies on the presence of DNA (Figure 5E). Only a small fraction of CcrM co-immunoprecipitated with LonS674A in the absence of DNA under the physiological concentrations of CcrM (1 µM) (Figure 5E). When we performed similar assays using elevated concentrations of the CcrM substrate (4 µM), CcrM co-immunoprecipitated with LonS674A, independent of the presence of DNA (Figure 5E). These results support our suggestion that DNA-dependent CcrM recognition by Lon occurs under physiological concentrations of CcrM. We determined that the intracellular concentration of CcrM is approximately 1 µM (Figure S3D). The *in vitro* degradation of CcrM by Lon is dependent on DNA when CcrM is present at 0.4 µM, but not at 4 µM CcrM, which is 4 times higher than the physiological concentration. Taken together, we propose a model where the robust degradation of CcrM requires both CcrM and Lon to interact while bound to DNA during the processive movement of CcrM (Figure 5F). The binding of the Lon protease to DNA does not allosterically stimulate substrate degradation. Instead, DNA plays an adaptor role in facilitating Lon recognition of CcrM under physiological conditions.

### Dynamic sequestration of CcrM at the new pole discriminates *Caulobacter* swarmer and stalked cell cycles

To determine if CcrM is protected from interaction with its substrate DNA prior to its complete digestion by Lon, we imaged cells in which CcrM was tagged with YFP. We constructed a strain expressing a sole chromosomal copy of *ccrM, yfp-ccrM*, under the control of its native promotor. YFP-CcrM fully complemented a *ΔccrM* strain. Strikingly, fluorescence microscopy revealed that YFP-CcrM formed a focus at the pole opposite the SpmX stalked pole marker (Jiang et al., 2014; Perez et al., 2017), demonstrating that YFP-CcrM accumulated at the new cell pole of the progeny stalked cell generated by cell division (Figure 6A). Among 444 analyzed cells in a mixed population, we observed that 29.50% of cells (n = 131) had a unipolar focus, while 40.99% of cells (n = 182) showed no detectable florescent signal. We also observed diffuse signal in 18.92% of examined cells (n = 84), suggesting that polar localization of CcrM is dynamic (Figure 6A). In these experiments, the cell population had two different types of stalked cells; those that result from the swarmer-to-stalked cell transition (that do not contain CcrM, Figure 2A) and those that result from cell division, accounting for the large population of cells with no fluorescent signal.

**Figure 6.**
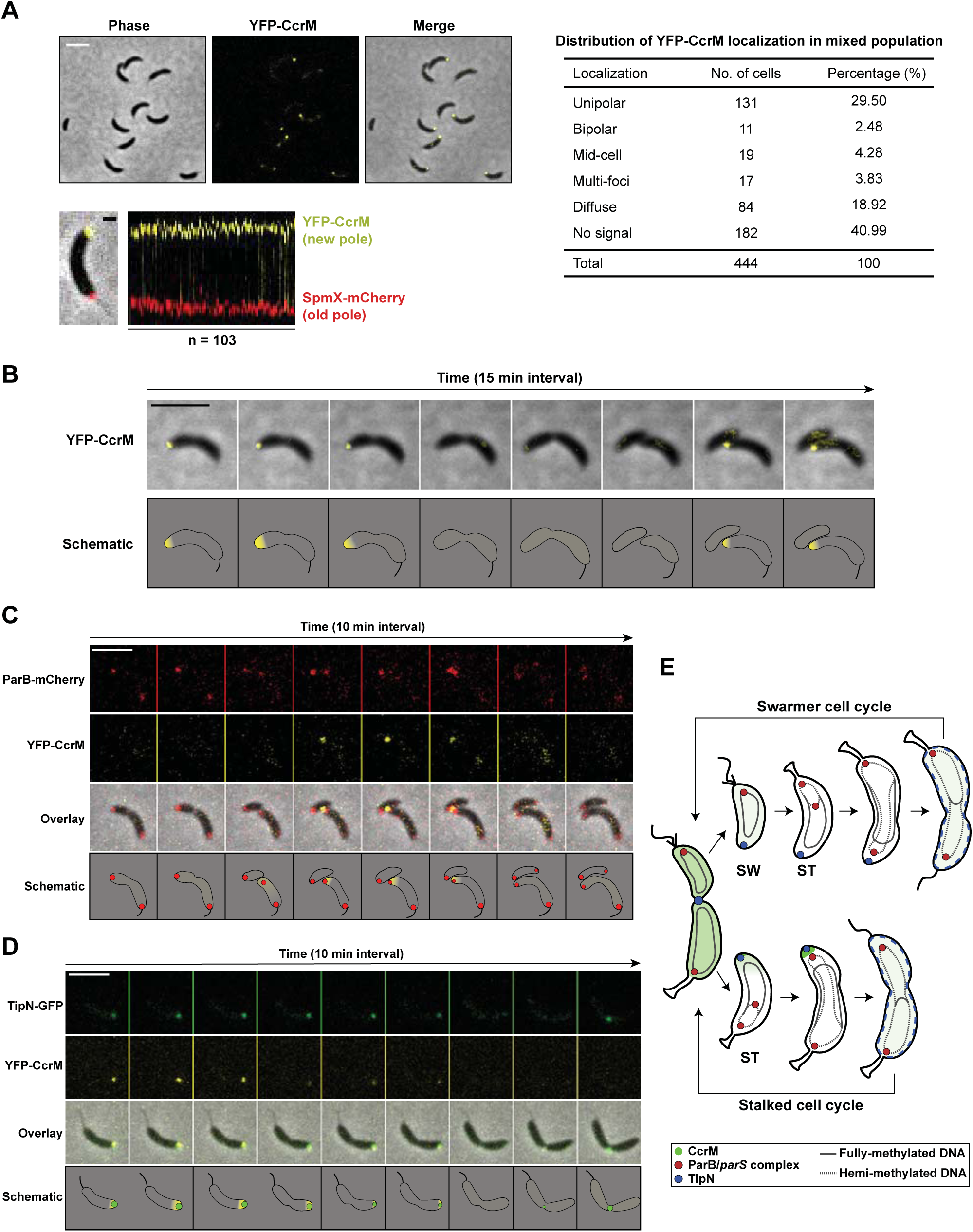
CcrM is dynamically sequestered at the flagellated cell pole of the stalked cell during stalked cell cycle. (A) Cells expressing single chromosomal copy of YFP-CcrM under the control of CcrM native promotor were grown in M2G to exponential phase and imaged by phase contrast and epifluorescence microscopy (upper left, Scale bar = 5 µm). Cells co-expressing YFP-CcrM and SpmX-mCherry under the control of their native promotors were grown in M2G to exponential phase and imaged by phase contrast and epifluorescence microscopy (lower left). A representative cell overlaid with phase, YFP, and mCherry channels is shown (Scale bar = 1 µm). A florescent profile is shown by an alignment of 103 cells with their fluorescent channels of pole marker SpmX-mCherry and YFP-CcrM. The table shows the distribution of CcrM localizations in examined 444 cells (right). (B) Time-lapse microscopy of cells producing chromosome-encoded YFP-CcrM under the control of its native promotor. Images of the cells were taken every 15 min. Scale bar = 5 µm. (C) and (D) Time-lapse microscopy of cells co-expressing chromosome-encoded YFP-CcrM and TipN-GFP (D) or ParB-mCheery (E) under the control of their native promotors. Images of the cells were taken every 10 min. Scale bar = 5 µm. (E) Cartoon depicting dynamic distribution of CcrM between swarmer and stalked cell cycle. CcrM protein abundance reaches the highest level in pre-divisional cell. Upon cell division, Swarmer (SW) daughter cell is subjected to the developmental program while stalked (ST) daughter cell begins chromosomal replication and cell growth immediately, giving raise to distinct swarmer and stalked cell cycle. In daughter swarmer cell, CcrM is degraded completely during swarmer to stalked cell transition. In daughter stalked cell, however, newly synthesized chromosomal DNA requires robust clearance of CcrM protein to maintain its hemi-methylated state, which cannot be achieved by proteolysis in a short time window. CcrM starts sequestration at the new pole when chromosomal replication initiated. Sequestered CcrM releases from the pole prior to the formation of division plane, meanwhile TipN is re-localized to the membrane throughout the cell.

To examine the subcellular distribution of CcrM during the cell cycle originating from the stalked cell arising from a cell division (Figure 6E), we used time-lapse microscopy to track cells (n > 100) that had a YFP-CcrM florescent focus at the new pole. A YFP-CcrM focus was consistently detected at the new pole of the progeny stalked cell and faded away during the transition to a pre-divisional cell (Figure 6B). Upon cell division, the YFP-CcrM focus appeared again at the incipient new pole of the stalked progeny cell, while no detectable signal was observed in the swarmer progeny (Figure 6B). To obtain the precise time of CcrM’s polar presence according to cell cycle milestone events, we carried out time-lapse microscopy of cells co-expressing YFP-CcrM and ParB-mCherry or TipN-GFP. ParB is a DNA-partitioning protein that binds to the centromeric *parS* locus near the chromosomal origin of replication. Localization of ParB reflects the movement of the ParB-bound centromere from the old pole to the new pole immediately upon the initiation of DNA replication (Ptacin et al., 2010). We observed the co-appearance of the YFP-CcrM focus and the ParB-*parS* complex at the new pole of the progeny stalked cell, suggesting that the sequestration of CcrM and initiation of chromosome replication begins at the same time (Figure 6C). In addition, TipN is a new cell pole marker that orients the polarity axis and its medial relocation reflects Z-ring formation at the division plane (Huitema et al., 2006; Lam et al., 2006). YFP-CcrM co-localized with TipN-GFP at the new pole of stalked cells (Figure 6D). During the transition from the stalked cell to the pre-divisional cell, when TipN-GFP left the new pole and started relocating to mid cell, YFP-CcrM was released from the pole together with TipN-GFP, demonstrating that CcrM polar sequestration ends prior to the formation of the division plane (Figure 6D). We did not observe an interaction between CcrM and TipN in a bacterial two-hybrid assay, implying that releasing of CcrM from the cell pole might be independent of the release of TipN (Figure S5A). We propose that any CcrM not cleared from the cell by proteolysis in the short time between cell division and the initiation of replication in the progeny stalked cell is inactivated by sequestration, thereby enabling the activation of gene transcription that requires hemi-methylated promotors. CcrM is dynamically sequestered to the new pole of only the stalked cell progeny. Thus, the distinct CcrM localization pattern between the two progeny cells thus discriminates *Caulobacter* swarmer and stalked cell cycles (Figure 6E).

### Polar sequestration stabilizes CcrM by preventing physical contact with Lon on DNA

Given that dynamic sequestration of CcrM at the new pole of the stalked cell occurs during the stalked cell cycle, and robust clearance of CcrM requires DNA as an adaptor, it is tempting to speculate that there are distinct patterns of CcrM proteolysis during the swarmer and stalked cell cycles (Figure 6E). Accordingly, we isolated progeny swarmer and stalked cells generated after the division of pre-divisional cells obtained from a synchronized cell population. Each progeny cell was then allowed to proceed through the cell cycle until 160 mps (Figure 7A II). CcrM in the progeny swarmer cell was degraded within 20 min (Note that in Figure 7A I, the swarmer cell population obtained from the original synchrony of a mixed population of cells contains swarmer cells primarily from 20-30 min of their development, by which time CcrM is completely degraded). CcrM was not completely cleared from the progeny stalked cell (Figure 7A II) although it was completely cleared from the stalked cell that resulted from the swarmer to stalked cell transition (Figure 7A I). As the progeny stalked cell progressed to the pre-divisional cell, we observed an increased abundance of CcrM commensurate with the increased synthesis of CcrM. (Figure 7A II). *In vivo* stability assays revealed that CcrM was quite stable although Lon was present (Figure S6A). We also observed a greater stability of CcrM in mixed swarmer and stalked progeny cells (collected at 160 mps) than in pre-divisional cells (collected at 120 mps) after 10 min shutoff of protein synthesis (Figure S6B). In *Δlon* cells, CcrM protein levels were stable throughout the experiment (Figure S6B).

**Figure 7.**
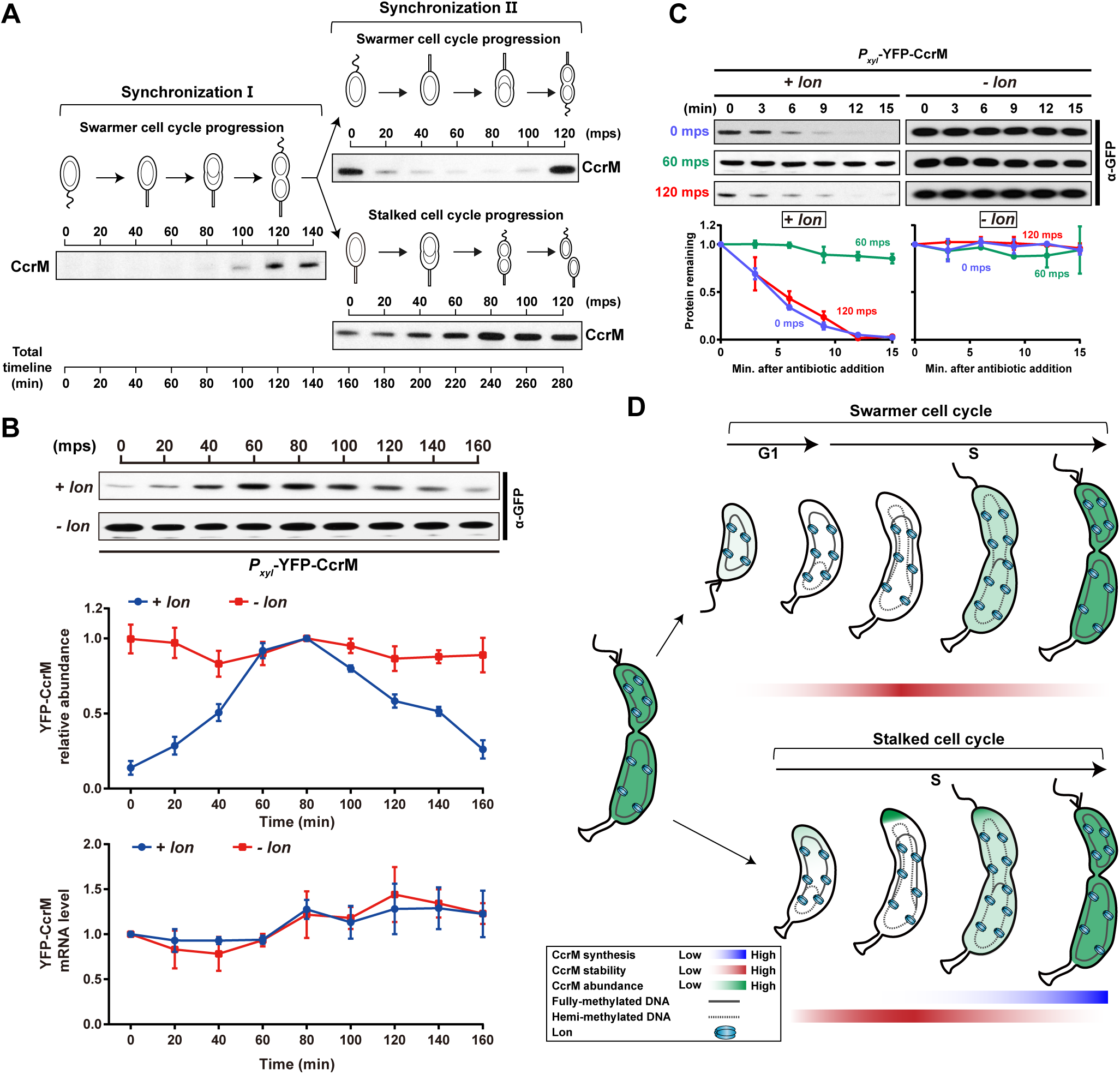
CcrM is stabilized by polar sequestration. (A) Immunoblots of protein samples from synchronized wild-type cultures using anti-CcrM antibody. Swarmer cells collected from the first synchronization were released into M2G medium allowing for cell cycle progression, harvested at 160 mps, and subjected to the second-synchronization. The swarmer and stalked cell fractions collected from the second synchronization were released in M2G for swarmer and stalked cell cycle analyses, respectively. (B) Immunoblots of protein samples from synchronized cells expressing YFP-CcrM under the control of *P*_*xyl*_. Merodiploid strains expressing YFP-CcrM in the background of wild-type (*+ lon*) or ∆*lon* (*- lon*) were grown in M2G with 0.3% xylose, synchronized and released into M2G with 0.3% xylose for cell cycle progression. Samples were taken every 20 min and protein levels were monitored by immunoblot using anti-GFP antibody (top). Band intensities were quantified (middle) and error bars represent SDs (*n* = 3). YFP-CcrM mRNA levels from each sample were normalized by qRT-PCR (bottom). The means ± SDs (*n* = 3) are plotted. (C) *In vivo* degradation assays showing YFP-CcrM stabilities in swarmer, stalked, and pre-divisional cell. YFP-CcrM stabilities in ∆*lon* (*- lon*) are shown for comparison. Merodiploid strains expressing YFP-CcrM controlled by *P*_*xyl*_ were grown in M2G with 0.3% xylose, synchronized, and harvested at 0, 60, and 120 mps. Samples were treated with antibiotics for protein synthesis shut-off assays. Protein levels were monitored by immunoblot using anti-GFP antibody (top). Band intensities were quantified (bottom) and error bars represent SDs (*n* = 3). (D) Cartoon depicting CcrM protein synthesis, stability and abundance between swarmer and stalked cell cycle. CcrM protein level reaches the highest point in pre-divisional cell. Meanwhile, CcrM starts proteolysis by Lon in a DNA-facilitated manner. Upon cell division, distinct CcrM protein turnover discriminates swarmer and stalked cell cycle. In swarmer cell cycle, remaining CcrM inherited from pre-divisional cell is completely degraded during swarmer-stalked cell transition (G1) via DNA-facilitated proteolysis. The transcription and translation of CcrM are repressed in early S phase and re-activated in late S phase. Although CcrM can be stabilized in early S phase, the protein abundance reaches its lowest point due to repressed transcription and translation. In stalked cell cycle, remaining CcrM inherited from pre-divisional cell is sequestered at the flagellated cell pole, which allows the initiation of chromosome replication at the stalked pole. The sequestration stabilizes CcrM during S phase by preventing physical contact with protease Lon.

The stable presence of CcrM in stalked cells derived from pre-divisional cells (Figure 7A II) suggests that CcrM may be specifically protected from Lon proteolysis by sequestration away from its DNA target during the stalked cell phase. To determine how CcrM degradation rate changes as a function of cell cycle, we created CcrM merodiploid strains by inserting *yfp-ccrM* under the control of the inducible xylose promoter as a single copy on the chromosome in wild-type or Δ*lon* backgrounds. In this genetic background, which equally produces CcrM at all cell cycle phases rather than only during pre-divisional cells, changes in protein abundance reflect changes in degradation rate. We performed immunoblots to monitor the presence of YFP-CcrM throughout the swarmer cell cycle. Interestingly, we found that in merodiploid cells containing Lon, the YFP-CcrM levels were low in swarmer cells, increased to the highest amount at ~ 80 mps, and decreased again during later stages of the cell cycle (Figure 7B). In contrast, the YFP-CcrM levels were constant during cell cycle progression in the *Δlon* background. Thus, CcrM is protected from Lon degradation between 60 mps and 80 mps of the swarmer cell cycle, which is consistent with the timing of CcrM sequestration during the stalked cell cycle.

To confirm differential CcrM turnover when CcrM is constitutively present during the cell cycle, we measured YFP-CcrM stabilities in merodiploid cells in the presence and absence of Lon. Translation shutoff assays by antibiotic addition were carried out using samples collected at 0 mps, 60 mps, and 120 mps during the swarmer cell cycle. We observed a robust degradation of YFP-CcrM protein in samples taken at 0 and 120 mps with measured half-lives of ~7 min in the presence of Lon (Figure 7C). For cells grown in the presence of Lon that were collected at 60 mps, however, YFP-CcrM degradation was not observed (Figure 7C). As a control, degradation was not observed in the absence of Lon at all time points.

In conclusion, CcrM is proteolyzed in swarmer cells, and there is no CcrM present in these cells until it is resynthesized during the pre-divisional stage. Though Lon is active during all phases of the cell cycle (Figure 3A), stalked cells can specifically sequester any CcrM that is present at the new cell pole, protecting it from interaction with DNA and the proteolysis by DNA-bound Lon during this phase (Figure 6B, Figure 7C). Thus, the CcrM inherited by stalked progeny from a pre-divisional cell is kept in an inactive state and consequently protected from proteolysis (Figure 7D). More broadly, this finding indicates that stalked cells arising from pre-divisional cells fundamentally differ from swarmer-derived stalked cells in their protein content, thus representing two distinct variants of the cell cycle.

## Discussion

Here we propose a model of CcrM protein turnover determined by coordinated DNA-facilitated protein degradation and CcrM sequestration during cell cycle progression. Although Lon is present and active throughout the cell cycle, CcrM transcription and translation is confined to the pre-divisional cell (Schrader et al., 2016; Zhou et al., 2015). At this time in the cell cycle, CcrM wins the race between synthesis and degradation and CcrM proceeds to processively methylate GANTC sites on the chromosome (Kozdon et al., 2013; Woodcock et al., 2017). When CcrM synthesis stops, Lon continues to clear CcrM from the cell. We show that robust degradation of CcrM by Lon requires the presence of DNA as an adaptor. Both CcrM and Lon have ~14-fold higher affinities for DNA than for each other, contributing to the high efficiency of DNA methylation of ~4500 GANTC sites by only ~600 CcrM molecules during a short time window of the cell cycle. Upon cell division, CcrM protein turnover varies between two daughter cells, giving rise to distinct swarmer and stalked cell cycles (Figure 7D). In the swarmer cell cycle, remaining CcrM inherited from pre-divisional cell is completely degraded during the swarmer-stalked cell transition (G1) via DNA-facilitated proteolysis (Figure 7D). Transcription and translation of CcrM are repressed in early S phase and re-activated in late S phase. The abundance of CcrM reaches its lowest point in early S phase due to repressed transcription and translation. In the stalked cell cycle, remaining CcrM inherited from the pre-divisional cells is sequestered to the new cell pole, concurrent with the immediate initiation of chromosome replication at the stalked cell progeny (Figure 7D). This sequestration of CcrM prevents DNA re-methylation during replication while also preventing its degradation by eliminating physical contact with the DNA-bound protease Lon. The sequestered CcrM is released from the pole at the time of new CcrM synthesis in the pre-divisional cell.

### DNA facilitated-proteolysis verses allosteric stimulation by other Lon substrates or unfolded proteins

Compared to the ClpXP protease that utilizes diverse adaptors for substrate delivery, Lon protease appears to process its substrate by directly recognizing clusters of exposed hydrophobic residues within a given polypeptide with little sequence specificity (Gur and Sauer, 2008). The first Lon substrate-specific adaptor, SmiA (swarming motility inhibitor A), was recently identified in *Bacillus subtilis*. Lon degrades the master flagellar activator protein SwrA only in the presence of SmiA. SmiA-dependent proteolysis is abolished upon surface contact causing SwrA protein levels to be stabilized and consequently increase motility (Mukherjee et al., 2015). In *Caulobacter*, Lon has been shown to degrade DnaA under proteotoxic stress leading to a cell cycle arrest (Jonas et al., 2013). *In vitro* experiments demonstrated that Lon alone cannot robustly degrade DnaA, but the addition of an unfolded substrate can allosterically activate Lon (Jonas et al., 2013). Similarly, heat shock protein Q (HspQ) was identified as a unique specificity-enhancing factor of Lon (Puri and Karzai, 2017). The addition of HspQ allosterically activates Lon and enhances the degradation of YmoA, a small histone-like protein whose efficient removal is required for bacterial virulence (Puri and Karzai, 2017). Given that adaptor-mediated proteolytic specificity for Lon protease is quite varied, Lon may employ multiple distinct mechanisms to regulate substrate specificity and degradation.

DNA binding activity of Lon was discovered three decades ago (Charette et al., 1984). Although several lines of evidence suggested that Lon binding to DNA can stimulate its ATPase activity and substrate degradation, the roles of this interaction in regulating substrate specificity and degradation remained to be elucidated. We showed here that the robust degradation of CcrM requires the binding of substrate and protease to DNA (Figure 4 and 5). Notably, DNA mediated activation of Lon degradation of CcrM cannot be ascribed to stimulated ATPase activity upon binding to DNA. The presence of substrate alone can induce the ATPase activity to a level similar to that induced by the co-presence of substrate and DNA (Figure 5C) and the presence of DNA does not stimulate degradation of non-DNA binding substrates (Figure S4A). Our results demonstrate that DNA serves as an adaptor for Lon-mediated CcrM proteolysis by facilitating substrate recognition rather than allosterically regulating Lon proteolytic activity. As a substrate for CcrM, DNA moonlights as an adaptor aiding CcrM delivery to the protease, which also prevents early degradation of CcrM prior to chromosomal methylation. In mitochondria of eukaryotic cells, Lon mutations were shown to be involved in multiple genetic diseases and cancer (Pinti et al., 2016). In prokaryotes, Lon is known to degrade multiple transcriptional regulators controlling the cell cycle, biofilm formation, motility and stress tolerance, and virulence (Breidenstein et al., 2012; Matsui et al., 2003; Rogers et al., 2016; Wright et al., 1996). Examples of Lon substrates in *Caulobacter* include CcrM, SciP and DnaA (Gora et al., 2013; Jonas et al., 2013; Wright et al., 1996), which all contribute to cell cycle regulation by their DNA binding activities. It is therefore conceivable that DNA-facilitated proteolysis may be a universal regulatory mechanism for specific recognition and degradation of DNA binding substrates. A corollary to this model is that Lon could temporally degrade a given substrate based on its own DNA-binding characteristics, so that a degradation hierarchy can be accommodated by a single factor and be achieved on a single platform. However, DNA-binding substrates other than CcrM, which lack processive movement along the DNA, may be regulated in a different mode or require involvement of other accessory factors.

### CcrM sequestration to the new cell pole

Bacterial cells employ multiple mechanisms to drive protein localization to the cell poles (Laloux et al., 2014; Rudner and Losick, 2010). We observed that the CcrM DNA methyltransferase is dynamically sequestered to the new pole of the progeny stalked cell (Figure 6). *Caulobacter* has been shown to recruit proteins to the cell poles through interaction with proteins or protein complexes that are already positioned at the pole. For example, the polar PopZ protein forms a microdomain that anchors the chromosome origin via its interaction with the chromosome partition complex ParB-*parS* (Bowman et al., 2008; Ebersbach et al., 2008). In addition, the stalked pole-localized protein, SpmX, serves as a bridge to direct the interaction between the DivJ histidine kinase and PopZ microdomain (Perez et al., 2017). Although the mechanism that localizes PopZ to the pole is not known, the PopZ microdomain captures multiple signaling proteins, thereby integrating several cellular processes within this membranes-less organelle (Bergé and Viollier, 2017; Holmes et al., 2016; Lasker et al., 2017). However, CcrM polar foci were observed in *ΔpopZ* strains, arguing that CcrM sequestration is PopZ-independent (Figure S5B). Assays of CcrM polar localization in strains lacking new pole-located proteins, including *ΔmopJ, ΔpodJ, and* a truncated *divL* (*divLΔ28*) showed that these proteins were also not necessary for CcrM sequestration (Figure S5B). A bacterial two-hybrid assay showed that CcrM does not interact with PleC, TipN, nor TipF (Figure S5A). Although unlikely that these proteins play a role in polar sequestration of CcrM, it is possible that CcrM can be captured by as yet unknown proteins so that the remaining CcrM molecules are not free to bind chromosomal DNA.

Mechanisms other than protein interaction may enable CcrM polar sequestration. Both CcrM and TipN are released from the cell pole prior to the formation of division plane (Figure 6D). The signals that trigger the dissociation of CcrM from the new pole are unknown. Narayanan and colleagues reported dynamic intracellular redox rhythms during the *Caulobacter* cell cycle (Narayanan et al., 2015), which precisely correspond to the dynamics of CcrM sequestration. The cytoplasm of the swarmer cell is in a reduced state during the G1 phase of the cell cycle. The reduced state then shifts to an oxidized state during the swarmer-to-stalk transition and early S phase. In late S phase, the stalked compartment of the pre-divisional cell remains in an oxidized state, while the swarmer compartment enters a reduced state. Intracellular redox state controls protein function and localization through formations of cysteine disulfide bond (Cremers and Jakob, 2013; Mou et al., 2003). In *Caulobacter*, NstA, a negative switch for topoisomerase IV (topo IV), inhibits decatenation activity of the topo IV by binding to the ParC DNA-binding subunit of topo IV (Narayanan et al., 2015). The activation of NstA requires dimerization by formation of intermolecular cysteine disulfide bonds under oxidizing conditions in early S phase. Trx1 was recently reported to be specifically induced in early S phase to counteract oxidizing stress (Goemans et al., 2018). CcrM contains two conserved cysteines (C13 and C329), of which the latter is located in a motif that is critical for DNA binding activity. It is possible that the localization of CcrM could be regulated by redox changes during the cell cycle.

### Asymmetric sequestration of CcrM fine tunes the access of CcrM to its DNA substrate

Replication is initiated on a fully methylated chromosome in the progeny stalked cell and in the stalked cell that arises from the swarmer-to-stalked cell transition. We have provided evidence that CcrM is processed differently in these two types of stalked cells. The progeny swarmer cell, which cannot initiate DNA replication, has 1/3 of the cell cycle to clear out CcrM before it differentiates into a stalked cell and its concurrent initiation of replication. The progeny stalked cell, on the other hand, immediately initiates replication and has very little time to clear out remaining CcrM. We have discovered that *Caulobacter* has devised a way to sequester CcrM so that it is not available to methylate the newly replicated strands of DNA in the progeny stalked cell, which would compromise cell cycle progression.

To prevent any residual CcrM activity, we observed that CcrM is sequestered to the new cell pole (Figure 6) where we hypothesize that it is prevented from accessing DNA. In support of this, we observed that sequestered CcrM is not degraded by DNA-bound Lon. Further, it is likely that CcrM does not bind chromosomal DNA at or near the origin of replication when sequestered at the pole because the ParB-*parS* complex is dissociated from the cell poles in *ΔpopZ* strain (Bowman et al., 2008; Ebersbach et al., 2008), while CcrM remains at the pole in a PopZ deletion strain (Figure S5B). The sequestration of CcrM would allow the newly synthesized chromosomal DNA to remain in the hemi-methylated state, thereby maintaining temporal control of transcription of cell cycle-regulated genes as the function of the passage of the replication fork. On the other hand, polar sequestration of CcrM would also prevent physical interaction with the DNA-bound Lon protease, thus stabilizing sequestered CcrM during S phase in the stalked cell cycle (Figure 7B). We propose that in the pre-divisional cell of the stalked cell cycle, sequestered CcrM is released from the pole, where it and newly synthesized CcrM binds DNA for processive m6A catalysis. The different patterns of CcrM degradation and sequestration during the swarmer and stalked cell cycles provide a fine-tuning mechanism that ensures that the immediate chromosome replication in the progeny stalked cell can proceed in the absence of re-methylation during DNA replication.

## Methods

### Bacterial strains, plasmids and growth conditions

Bacterial strains and plasmids used in this study are listed in Table S1. Primers used for this study are listed in Table S2. *E. coli* strains were routinely grown in LB medium at 37 °C with appropriate antibiotics (100 µg ml^−1^ ampicillin, 50 µg ml^−1^ kanamycin). *Caulobacter* strains were grown in PYE (rich medium) or M2G (minimal medium) at 37°C, supplemented with 0.3% xylose when necessary. Antibiotics were supplemented as needed for solid and liquid media, respectively, with the following concentration: kanamycin (25 µg ml^−1^ or 5 µg ml^−1^), spectinomycin (50 µg ml^−1^ or 25 µg ml^−1^), oxytetracycline (2 µg ml^−1^ or 1 µg ml^−1^), gentamycin (10 µg ml^−1^ or 5 µg ml^−1^).

### Strain construction

To construct XZC13 and XZC14, plasmids pNP138-M2-CcrM and pNP138-CcrM-M2 were introduced into NA1000 by electroporation, respectively. Clones that have integrated the vector at the *ccrM* locus were selected on PYE plates containing kanamycin. A second recombination step was performed to select for plasmid excision. Colonies arising from the first integrants were grown in PYE plain for at least 6 hours. Cells were serious diluted for counter-selection on PYE containing 3% sucrose. Colonies grown on PYE sucrose plates were replicated on PYE containing kanamycin for selection of plasmid excision. Colonies that were able to grow on PYE sucrose, but not on PYE kanamycin plates were grown in liquid PYE plain medium for PCR verification.

To characterize the role of CcrM C-terminus in proteolysis, XZC34, XZC35, and XZC36 were constructed by electroplating pXYFPN2-CcrM, pXYFPN2-CcrM65C, and pXYFPN2-CcrMΔC65 into NA1000, respectively. XZC154, XZC161, and XZC88 were constructed by electroplating pXYFPN2-CcrM, pXYFPN2-CcrM65C, and pXYFPN2-CcrMΔC65 into LS2382, respectively.

To identify CcrM degradation tag, strains XZC105 and XZC160 were generated by electroporating pXYFPN2-CcrM30C into NA1000 or LS2382, respectively. Strains XZC139, XZC138, XZC144, XZC143, XZC137, XZC106, XZC108, XZC109, XZC114 were constructed similarly to XZC105, except that plasmid pXYFPN2-CcrM28C, pXYFPN2-CcrM26C, pXYFPN2-CcrM24C, pXYFPN2-CcrM23C, pXYFPN2-CcrM22C, pXYFPN2-CcrM20C, pXYFPN2-CcrM15C, pXYFPN2-CcrM10C, or pXYFPN2-CcrM8C, respectively, was used for electroporation. Strains XZC159, XZC158, XZC162, XZC163, XZC157, XZC164, XZC156, XZC155, XZC165 were constructed similarly to XZC160, except that plasmid pXYFPN2-CcrM28C, pXYFPN2-CcrM26C, pXYFPN2-CcrM24C, pXYFPN2-CcrM23C, pXYFPN2-CcrM22C, pXYFPN2-CcrM20C, pXYFPN2-CcrM15C, pXYFPN2-CcrM10C, or pXYFPN2-CcrM8C, respectively, was used for electroporation.

To mutate the conserved amino acids at C-terminal of CcrM, strains XZC121, XZC129, XZC135, XZC134, XZC131 were constructed by electroporating plasmid pXMCS2-CcrMD304A, pXMCS2-CcrMS315A, pXMCS2-CcrMW332A, pXMCS2-CcrMS350A or pXMCS2-CcrMS347A into NA1000, respectively. The integration of the plasmid at the *ccrM* locus was further verified by PCR.

To identify Lon activity during cell cycle, strains XZC6 and XZC86 were generated by electroporating pXYFPN2-sul20C into NA1000 or LS2382, respectively. To investigate the subcellular localization of Lon protease, strains XZC142 and XZC148 were generated by electroporating plasmid pXYFPC2-Lon or pXYFPC2-LonQM into PC6340, respectively. Strains XZC20 was generated by electroporating plasmid pCHYC2-Lon into NA1000. XZC23 was constructed similarly to XZC13, except that the plasmid pNP138-mCherry-Lon was used for electroporation.

To identify the dynamic localization of CcrM during cell cycle, XZC24 was constructed similarly to XZC13, except that the plasmids pNP138-YFP-CcrM was used for electroporation. Strains XZC75 and XZC112 were generated by electroporating plasmid pCHYC1-SpmX or pCHYC1-ParB into XZC24, respectively. XZC89 was constructed by transducing *tipN-gfp (gent^r^)* from CJW1406 into XZC13.

To observe whether CcrM localization is dependent on polar localized proteins, strain XZC49 was constructed by phage transducing from GB255 into XZC24. XZC50 was constructed by electroporating pXYFPN2-CcrM into LS4461. Strains XZC68, XZC71, XZC69, XZC70 were constructed by electroporating plasmid pMCS2-podJ, pMCS2-mopJ, pMCS2-perP or pMCS2-spmX into XZC24, respectively.

### Expression plasmids

To generate pET28b-CcrM, the *ccrM* ORF including stop codon was amplified using KOD DNA Polymerase (EMD Millipore) and inserted into pET28b digested with *Nde*I and *Eco*RI via Gibson assembly (NEB). Plasmid pET28b-Lon was generated similarly to pET28b-CcrM, except PCR amplification of the *lon* ORF. Plasmids pET28b-CcrMΔC65, pET28b-CcrMS315A, pET28b-LonS674A, and pET28b-LonQM were generated by mutagenesis using Q5 Site-Directed Mutagenesis Kit (NEB).

### Integrating plasmids

The integration vector pNP138-M2-CcrM was constructed by amplifying an upstream and downstream homology region of *ccrM* using primer pairs M2ccrmLB-F/M2ccrmLB-R and M2ccrmRB-F/M2ccrmRB-R, respectively. The two fragments were inserted into *Spe*I-*Eco*RI digested vector pNPTS138 via Gibson assembly to yield pNP138-M2-CcrM. pNP138-CcrM-M2 was constructed similarly to pNP138-M2-CcrM, except that primer pairs ccrmM2LB-F/ccrmM2LB-R and ccrmM2RB-F/ccrmM2RB-R were used for PCR amplification. pNP138-YFP-CcrM and pNP138-mCherry-Lon were generated using a similar strategy.

To construct pXYFPN2-CcrM, the *ccrM* ORF was amplified and inserted into *Kpn*I-*Eco*RI digested pXYFPN2 via Gibson assembly. pXYFPN2-CcrMΔC65 was constructed similarly to pXYFPN2-CcrM, except amplification of *ccrM* ORF lacking C-terminal 65 amino acids. The other pXYFPN2-CcrM derivative plasmids were generated by Q5 mutagenesis using pXYFPN2-CcrM as the backbone. Primers used for mutagenesis are listed in Table S2.

To construct pXYFPN2-sul20C, primer pair sul20C-F/sul20C-R was used to amplify pXYFPN2 backbone and the sul20C degradation tag was inserted by Q5 mutagenesis. To construct pXYFPC2-Lon, the *lon* ORF lacking stop codon was amplified and inserted into *Nde*I-*Kpn*I digested pXYFPC2 via Gibson assembly. pXYFPC2-LonQM was generated by Q5 mutagenesis based on pXYFPC2-Lon using primer pairs Lon(4m)MU-F/Lon(4m)MU-R. To construct pCHYC2-Lon, the fragment encoding Lon 407-799 amino acids was amplified and inserted into *Nde*I-*Kpn*I digested pCHYC2 via Gibson assembly.

To construct pCHYC1-SpmX, the fragment encoding SpmX 207-431 amino acids was amplified and inserted into *Nde*I-*Kpn*I digested pCHYC1 via Gibson assembly.

To construct pCHYC1-ParB, the fragment encoding ParB 104-304 amino acids was amplified and inserted into *Nde*I-*Kpn*I digested pCHYC1 via Gibson assembly.

To construct pXMCS2-CcrM, the *ccrM* ORF was amplified and inserted into *Nde*I-*Kpn*I digested pXMCS2 via Gibson assembly. The resultant plasmid was used to generate pXMCS2-CcrMD304A, pXMCS2-CcrMS315A, pXMCS2-CcrMW332A, pXMCS2-CcrMR350A, and pXMCS2-CcrMD347A by Q5 mutagenesis.

To construct pKNT25-CcrM and pKT25-CcrM, the *ccrM* ORF was amplified and inserted into *Hin*dIII-*Bam*HI digested pKNT25 or *Bam*HI-*Eco*RI digested pKT25 via Gibson assembly, respectively. To construct pUT18-PleC, the fragment encoding PleC 7- 842 amino acids was amplified and inserted into *Hin*dIII-*Bam*HI digested pUT18 via Gibson assembly. To construct pUT18-PleC, the fragment encoding PleC 7-842 amino acids was amplified and inserted into *Hin*dIII-*Bam*HI digested pUT18 via Gibson assembly. To construct pUT18-DivL, the fragment encoding DivL 2-768 amino acids was amplified and inserted into *Hin*dIII-*Bam*HI digested pUT18 via Gibson assembly. To construct pUT18C-PodJ, the *podJ* ORF was amplified and inserted into *Bam*HI-*Eco*RI digested pUT18C via Gibson assembly. To construct pUT18-TipN, the fragment encoding TipN 2-882 amino acids was amplified and inserted into *Hin*dIII-*Bam*HI digested pUT18 via Gibson assembly. To construct pUT18-TipF, the fragment encoding TipF 2-452 amino acids was amplified and inserted into *Hin*dIII-*Bam*HI digested pUT18 via Gibson assembly.

To construct pMCS2-mopJ, the fragment encoding MopJ 6-144 amino acids was amplified and inserted into *Nde*I-*Nhe*I digested pMCS2 via Gibson assembly. To construct pMCS2-spmX, the fragment encoding SpmX 12-290 amino acids was amplified and inserted into *Nde*I-*Nhe*I digested pMCS2 via Gibson assembly. To construct pMCS2-podJ, the fragment encoding PodJ 28-423 amino acids was amplified and inserted into *Nde*I-*Nhe*I digested pMCS2 via Gibson assembly. To construct pMCS2-perP, the fragment encoding PerP 23-155 amino acids was amplified and inserted into *Nde*I-*Nhe*I digested pMCS2 via Gibson assembly.

### Caulobacter synchronization

The synchronization experiment was performed as previously described (Schrader and Shapiro, 2015). The synchronized swarmer cells were released into M2G medium supplied with certain antibiotics as needed. Samples were taken every 20 min for further analysis as indicated in the figure. For the double-synchronization experiment, the swarmer cells raised from the first synchronization were released and grown into M2G medium at 30 °C. Cells were collected at 160 minutes past synchrony (mps) and subjected to the second synchronization. The swarmer and stalked fractions were collected, released into M2G, and monitored for cell cycle progression every 20 min.

### Protein purification

*Caulobacter* Lon and its variants were purified using a combination of Ni-NTA affinity and size exclusion chromatography steps. ER2566 (NEB) harboring pET28b-Lon plasmid was grown in LB containing 50 µg/ml kanamycin and 3% ethanol, and protein expression was induced overnight at 16 °C with 1 mM IPTG at OD_600_ of 0.5. Cells were harvested and resuspended in purification buffer (50 mM HEPES pH 7.5, 100 mM NaCl, 100 mM KCl, 25mM imidazole, 10% Glycerol). After sonication, buffer-equilibrated Ni-NTA beads were added to cleared cell lysate, incubated at 4 °C for 1 hour, and washed extensively with purification buffer. The target protein was eluted with purification buffer containing 325 mM imidazole. The protein sample was buffer exchanged to column buffer (50 mM HEPES pH 7.5, 100 mM NaCl, 100 mM KCl, 2 mM β-ME), loaded on a Sephacryl S-200 column. Fractions containing Lon were pooled, concentrated, dialyzed against protein storage buffer (50 mM HEPES pH 7.5, 100 mM NaCl, 100 mM KCl, 10% Glycerol), and stored at −80 °C. CcrM and its variants were purified similarly to Lon. The removal of 6xHis tag was performed using Thrombin CleanCleave Kit (Sigma) and verified via immunoblot using anti-His antibody.

### Protein *in vivo* and *in vitro* degradation assays

For protein *in vivo* degradation assay, cells were grown under the desired conditions. Protein synthesis was blocked by addition of 200 µg/ml chloramphenicol and 1 mg/ml spectinomycin. Samples were taken at the time-points indicated in the figure and snap-frozen in liquid nitrogen before immunoblot analysis.

*In vitro* degradation assays were performed in Lon degradation buffer (100 mM KCl, 10 mM MgCl_2_, 1 mM DTT, and 25 mM Tris-HCl [pH 8.0]) at 30 °C with an ATP-regeneration system (10 U/ml rabbit muscle pyruvate kinase [or 75 µg/ml creatine kinase], 20 mM phosphoenolpyruvate [or 20 mM creatine phosphate], 4 mM ATP). The concentrations of Lon_6_, LonS674A_6_, LonQM_6_, CcrM, CcrMΔC65, CcrMS315A, or β-casein were 0.2 µM, 0.2 µM, 0.2 µM, 1 µM, 1 µM, 1 µM, and 1 µM respectively. Samples were taken every 30 min, quenched with SDS loading buffer, heated at 95 °C, and snap-frozen in liquid nitrogen. Samples were pre-warmed at 65 °C prior to separation by SDS-PAGE. The gels were stained by Coomassie blue G-250. Protein degradation rates were calculated based on quantification of protein band intensity using ImageJ.

CcrM remaining levels over reaction time were fit to a single exponential model equation

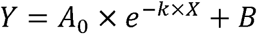

where Y is CcrM protein remaining, X is reaction time (min.), A_0_ is the initial amount of substrate (normalized to 1), *k* is degradation rate, and B is the fitting background. The fitting parameters over DNA concentrations were listed as follows:

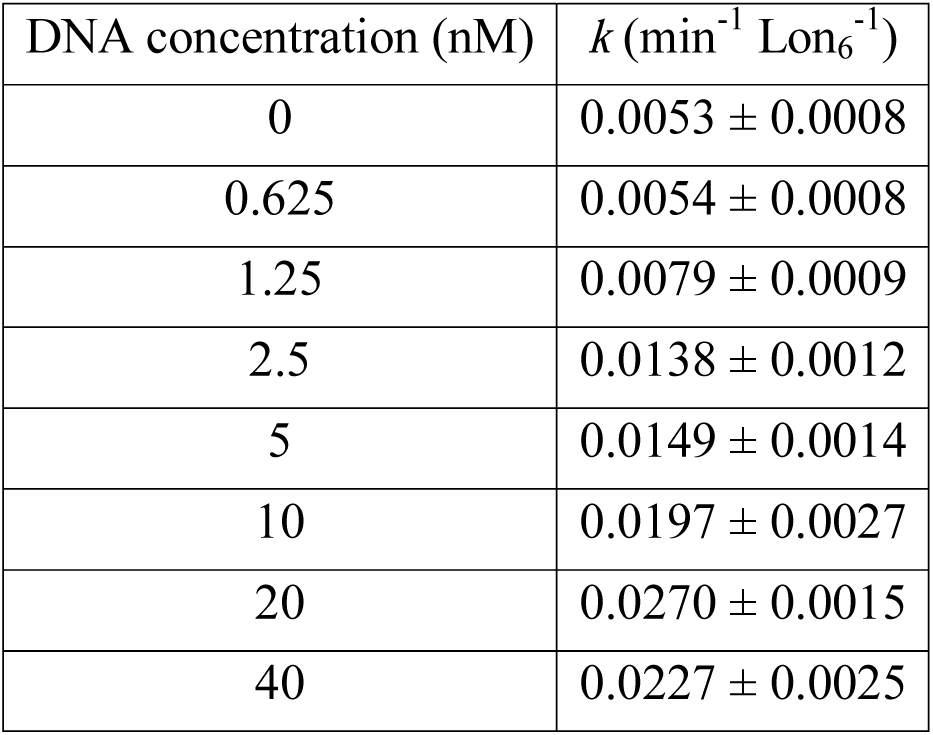

In Figure 5B, CcrM degradation rates over DNA concentrations were fit to an agonist-stimulated dose-response model:

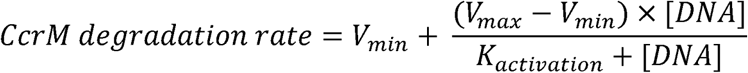

with fitted parameters: *V*_*min*_ = 0.0039 ± 0.0016, *V*_*max*_ = 0.0289 ± 0.0022 min^−1^ Lon_6_^−1^, *K*_activation_ = 4.391 ± 1.609 nM for DNA stimulation.

### Immunoblotting

Harvested cells were suspended in SDS loading buffer and heated for 10 min at 95 °C. Equal amounts of total protein were separate on 4–15% gradient polyacrylamide gel (Bio-Rad), semi-try transferred to PVDF membrane, and probed with appropriate dilutions of primary antibody against targeted protein indicated in the figure, and a 1: 10,000 dilution of secondary HRP-conjugated antibody. Washed membrane was developed using Super Signal West Pico Chemiluminescent Substrate (Thermo Scientific) and exposed to an X-ray film for visualization. The film was scanned, and the band intensity was quantified using ImageJ software.

### Quantitative reverse transcription PCR

Cells grown under the desired conditions were harvested, treated with two volumes of RNAprotect Bacteria Reagent (Qiagen), and snap-frozen in liquid nitrogen. The total RNA was extracted using the Qiagen RNeasy Mini Kit. Contaminated genomic DNA was removed through on-column digestion with a DNase using the Qiagen RNase-free DNase Kit. The RNA concentration was determined using a NanoDrop 2000 spectrophotometer (Thermo Scientific). Reverse transcription and cDNA synthesis were performed using QuantiTect Reverse Transcription Kit. Quantitative PCRs were performed using Luna Universal qPCR Master Mix (NEB) on an Applied Biosystems 7500 Fast Real-Time PCR system. The *rho* gene was used as an endogenous control. The relative fold change in target gene expression was calculated using a 2^−ΔΔCT^ method (Schmittgen and Livak, 2008).

### Microscopy

*C. crescentus* strains grown to exponential phase (OD_600_ < 0.3) and spotted on agarose pads (1.5%) containing M2G prior to imaging. Phase-contrast and fluorescence microscopy images were obtained using a Leica DMi8 microscope with an HC PL APO 100×/1.40 oil PH3 objective, Hamamatsu electron-multiplying charge-coupled device (EMCCD) C9100 camera, and Leica Application Suit X software. For all image panels, the brightness and contrast of the images were balanced with ImageJ (NIH) to represent foci or diffuse fluorescent signal. For computational image analyses, MicrobeJ (Ducret et al., 2016) was used to determine cell outlines and lengths from phase images. Oufti was used to determine normalized fluorescence intensities from each single cell. The data was plotted and statistically analyzed using Prism 7 (GraphPad).

### Measurement of fluorescence intensity in living cells

Cells grown under the desired condition were diluted to OD_600_ of 1. A 300 µl aliquot of cell suspension was added to the each well of a 96-well plate. The absolute fluorescence intensity was measured using Tecan Infinite M1000 plate reader at the High-Throughput Bioscience Center (HTBC), Stanford.

### *In vitro* DNA methylation and ATPase assays

Probe 1 was amplified from NA1000 genome using primer pair probe1-F/probe1-R. Probe 2 was generated by double-joint PCR, using Probe 1 as a template. Two resultant fragments amplified by primer pairs probe1-F/probe1-RMu and probe1-FMu/probe1-R were jointed using the second-round of PCR with primer pair probe1-F/probe1-R. Probe 3 was amplified from NA1000 genome using primer pair probe3-F2/probe3-R2. To prepare fully methylated DNA probe, 20 nM Probe 1 was incubated with 420 nM CcrM and 80 µM S-adenosyl-methionine (SAM) at 30 °C for 1 hour in DNA methylation buffer (50 mM Tris-HCl [pH 7.5], 5 mM β-ME, 10 mM EDTA). The resultant DNA probe was precipitated with 100% ethanol, washed twice with 70% ethanol, dried in speed-vac, and re-dissolved in distilled-water. To prepare hemi-methylated Probe 1, an equal amount of fully methylated and unmethylated Probe 1 was mixed in a PCR tube. The denature and annealing were performed on a thermocycler with 3 min at 95 °C following 3 min at 70 °C for 5 cycles. The resultant DNA probe was precipitated with 100% ethanol, dried in speed-vac, and re-dissolved in distilled-water. The methylation state of probe was assayed by restriction digestion using *Hinf*I or *Hph*I. Lon ATPase activity was assayed using ATPase/GTPase Activity Assay Kit (Sigma).

### Bacterial-two hybrid

The bacterial adenylate cyclase two-hybrid system was used to test protein interactions (Karimova et al., 1998). Briefly, genes of interest were fused to the N- or C-terminal of T18 or T25 fragments in the pUT18C pUT18C, pKT25, or pKNT25 vectors. The resultant plasmids were co-introduced into BTH101 strain. The transformants were re-streaked on MacConkey agar (40 g/L) plates supplemented with maltose (1%), IPTG (1 mM), and appropriate antibiotics. Plates were incubated at 30°C for 3 days before photography.

### Microscale thermophoresis (MST)

Fluorescent labeling of lysine residues in LonS674A was accomplished by incubating each protein with an N-hydroxysuccinimide (NHS) ester conjugated to Atto-488 (Sigma-Aldrich). The dye-conjugate was dissolved in dry DMSO to make a 1 mM solution. The conjugation reaction was performed in the dark using 1-2 mg/mL protein and a 3-fold molar ratio of dye to protein at room temperature, with gentle shaking. Unconjugated dye was removed through dialysis against the protein storage buffer. Direct binding between fluorescently labeled LonS674A and CcrM or Probe 1 was probed via microscale thermophoresis (NanoTemper Technologies) (Wienken et al., 2010). For each binding experiment, a twofold serial dilution was made for CcrM or Probe 1 in protein storage buffer with 0.025% Tween-20 and 10 mM MgCl_2_. Fluorescently labeled LonS674A was then added at 25 nM, mixed, and incubated at room temperature for 10 minutes, covered, in the dark. The protein mixtures were loaded into Standard Treated capillaries (NanoTemper). Binding was assessed using the following instrument settings: 70% blue LED power, 40% IR-laser power, 30 second IR heating period, 5 second recovery.

Binding data were initially fit in MO.Affinity Analysis (NanoTemper), and the binding curve plateau data were exported. Experimental replicates were averaged in Prism 7 (GraphPad) and according to the law of mass action, as described:

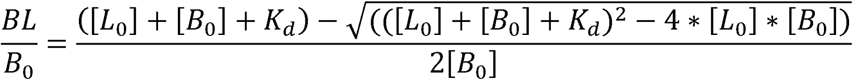

In this equation, BL represents the concentration of protein complexes, [*B*_0_] represents total binding sites of the fluorescent ligand, [*L*_0_] represents the amount of added ligand, and *K*_*d*_ represents the dissociation constant.

### Electrophoretic mobility shift assay (EMSA)

DNA binding capacity of CcrM was evaluated by incubation of purified CcrM with 20 nM of DNA probe indicated in the figure in the presence of 200 µM sinefungin in EMSA buffer (50 mM HEPES pH 7.0, 200 mM NaCl, 1 mM EDTA, 1 mM DTT) for 30 min at room temperature and subjected to electrophoresis in a 4–15% Mini-PROTEAN^®^ TGX^™^ Precast Protein Gels (Bio-Rad) at constant 80 V for 3 hours at 4°C in 1× Tris glycine native gel buffer (25 mM Tris base, 192 mM glycine). Lon DNA binding capacity was assayed similarly to CcrM, except that 10 mM MgCl_2_ was added instead of 200 µM sinefungin. Protein concentrations were 0 µM, 2 µM, 4 µM, 6 µM, 8 µM, 10 µM, 12 µM for CcrM and 0 nM, 50 nM, 100 nM, 150 nM, 200 nM, 250 nM for Lon_6_, respectively. The protein-DNA complexes were stained with ethidium bromide and imaged with a Bio-Rad ChemiDoc XRS+ system.

### *In vitro* Ni-NTA pull-down assay

Purified LonS674A_6_ (0.2 µM) was incubated with 20 nM Probe 1 and 200 µl buffer-equilibrated Ni-NTA beads at room temperature for 30 min in PD buffer (protein storage buffer containing 10 mM MgCl_2_). One unit of DNase I was added when necessary to cleavage Probe 1. The beads were washed once with 1 ml PD buffer and resuspended in another 200 µl PD buffer containing a low amount of CcrM (0.4 µM) or high amount of CcrM (4 µM). A 20 µl aliquot of reaction (input) was taken, suspended in SDS loading buffer, boiled for 10 min followed by incubation at 65 °C for 5 min, and subjected to analyses by SDS-PAGE and 1% agarose gel. The content of remaining reaction was incubated at room temperature for 1 hour, washed with PD buffer extensively, and eluted with 100 µl PD buffer containing 325 mM imidazole. The eluted protein samples were analyzed by SDS-PAGE for detection of the presences of LonS674A_6_-CcrM-DNA nucleoprotein complex via silver staining.

## Author contributions

X.Z. and L.S. initiated the study. X.Z. designed and performed experiments, performed data analysis. X.Z. and L.S. wrote the paper.

### Acknowledgement

We thank all members of the Shapiro lab for helpful discussions throughout the project, Drs. Thomas H. Mann, Jonathan Herrmann, Saumya Saurabh and Darshankumar T. Pathak for their critical reading of the manuscript, and Dr. Jinfan Wang for help on modelling protein degradation data. We acknowledge support from NIGMS NIH R35- GM118071 [to L.S.]. L.S. is a Chan Zuckerberg Biohub Investigator.

## Supplementary figure legends

**Figure S1. Conserved C-terminal motifs determine CcrM DNA binding activity, related to Figure 2**.

(A) Sequence alignment of CcrM homologs from twelve divergent α-proteobacterial species reveals four conserved motifs at C-terminus. The conserved residues subjected to mutation from each motif are highlighted.

(B) Phase contrast micrographs of *ccrM* depletion strains complemented with CcrM, CcrMD304A, CcrMS315A, CcrMW332A, CcrMD347A, CcrMR350A. Scale bar = 5 µm.

(C) Cell length analyses of strains in (A). Mean cell length (µm) ± SEM: CcrM = 3.10 ± 0.08 (*n* = 128); CcrMD304A = 2.71 ± 0.05 (*n* = 120); CcrMS315A = 7.66 ± 0.54 (*n* = 112); CcrMW332A = 9.57 ± 0.59 (*n* = 120); CcrMD347A = 2.78 ± 0.06 (*n* = 133); CcrMR350A = 2.92 ± 0.07 (*n* = 152). **** indicates *P* < 0.0001 by one-way ANOVA.

(D) Spot dilutions of strains in (A). Cells in exponential phase were diluted to an OD600 of 0.03, serially diluted and spotted onto the same PYE agar plate and incubated at 30 °C for 2 days before photography.

(E) EMSA showing abolished DNA binding activity caused by mutation at S315A on CcrM. See Methods for experimental details.

**Figure S2. Verification of Lon DNA-binding and proteolytic activities, related to Figure 3**.

(A) EMSA showing the effect of alanine substitutions at S674 and K301/K303/K305/K306 on Lon DNA binding activities. See Methods for experimental details.

(B) Phase contrast and epifluorescence images showing cell morphology and Lon distribution. Wild-type cells expressing chromosomal YFP-Lon or Lon-YFP under the control of native promotor were grown in M2G to exponential phase and imaged. Scale bar = 10 µm.

**Figure S3. Binding of CcrM to DNA is irrelevant to DNA methylation states, related to Figure 4**.

(A) Schematic view of restriction sites on Probe 1 and rationale of restriction digest-based DNA methylation assay. *Hinf*I is only able to cut unmethylated GANTC site (blue). *Hph*I cuts GGTGA(N)_8_ that overlapped with a half of GANTC site (brown). Adenine methylated GGTGA_m_(N)_8_ is resistant to *Hph*I digestion.

(B) Agarose gels showing the verification of DNA methylation states by restriction digest analyses with *Hinf*I and *Hph*I. Two DNA fragments are expected for *Hinf*I digestion

(C) EMSA showing CcrM and Lon binding to unmethylated, hemi-methylated and fully-methylated Probe 1. See Methods for experimental details.

(D) Quantitative immunoblots of CcrM levels in *Caulobacter* pre-divisional cell. Immunoblots were performed following SDS-PAGE of different concentrations of purified CcrM and a *Caulobacter* pre-divisional cell lysate collected at 120 mps. The intracellular concentration of CcrM was 1090 ± 135 nM or ∼ 600 ± 150 CcrM monomers per cell.

**Figure S4. DNA plays an adaptor role in CcrM proteolysis by Lon, related to Figure 5**.

(A) *In vitro* degradation assays showing the degradation of β-casein by Lon in the presence and absence of DNA. β-casein (1 µM) was incubated with Lon_6_ (0.2 µM) in the absence or presence of Probe 1 (10 nM). Creatine kinase is part of the ATP regeneration system.

(B) *In vitro* degradation assays showing the degradation of β-casein by LonQM. β-casein (1 µM) was incubated with LonQM_6_ (0.2 µM) in the absence or presence of ATP (4 mM). Creatine kinase is part of the ATP regeneration system.

**Figure S5. Identification of the roles of known polar localized proteins in CcrM sequestration, related to Figure 6**.

(A) Bacterial two-hybrid assays showing the negative interaction of CcrM to polar localized proteins (PleC, DivL, PodJ, TipN, and TipF). − / − and + / + indicate a negative and a positive control, respectively. Red colonies indicate a positive interaction. Cells were grown at 30 °C for 2 days before photography.

(B) Overlaid phase contrast and epifluorescence images showing CcrM polar sequestration in cells depleting several known polar localized proteins and protease regulator PerP. CcrM polar sequestration does not depend on the presence of DivL, PopZ, PodJ, MopJ, and PerP. Deletion of SpmX serves as a negative control.

**Figure S6. *In vivo* stability of CcrM, related to Figure 7**.

(A) *In vivo* degradation assays showing CcrM stability in a mixed population. Stabilities of YFP chimeric proteins in ∆*lon* (*- lon*) cells are shown for comparison. Cells were grown in PYE with 0.3% xylose to exponential phase and treated with antibiotics for protein synthesis shut-off assays. Protein levels were monitored by immunoblot using anti-GFP antibody (top). Band intensities were quantified (bottom) and error bars represent SDs (*n* = 3).

(B) *In vivo* degradation assays showing CcrM stabilities at 120 mps and 160 mps in wild-type. CcrM stabilities in ∆*lon* cells are shown for comparison. Cells were harvested at 160 mps or 120 mps and treated with antibiotics to shut-off protein synthesis. Protein levels were monitored by immunoblot using anti-CcrM antibody (top). Band intensities were quantified (bottom) and error bars represent SDs (*n* = 3).

